# PTGS is dispensable for the initiation of epigenetic silencing of an active transposon in *Arabidopsis*

**DOI:** 10.1101/2024.05.27.596030

**Authors:** Marieke Trasser, Grégoire Bohl-Viallefond, Verónica Barragán-Borrero, Laura Diezma-Navas, Lukas Loncsek, Magnus Nordborg, Arturo Marí-Ordóñez

## Abstract

Transposable elements (TEs) are largely repressed in plants through transcriptional gene silencing (TGS), which is maintained by heritable epigenetic silencing marks such as DNA methylation. However, the mechanisms by which silencing is installed in the first place remains poorly understood in plants. Small interfering (si)RNAs and post-transcriptional gene silencing (PTGS) play a role in the initial response by reducing mRNA and protein levels of active TEs and are believed to mediate the initiation of TGS by guiding the first deposition of DNA methylation. To determine how this silencing installation works, we took advantage of *ÉVADÉ (EVD)*, an endogenous retroelement in Arabidopsis, which can be used to recapitulate true *de novo* silencing with a well-established sequence of PTGS followed by a TGS phase. To test whether PTGS is a prerequisite for TGS, active *EVD* copies were introduced into RNA-DEPENDENT-RNA-POLYMERASE-6 (RDR6) mutants lacking an essential PTGS component. *EVD* activity and silencing were monitored across several generations. Unexpectedly, even in the absence of PTGS, TGS and silencing of *EVD* were still achieved through installation of RNA-directed DNA methylation (RdDM) at *EVD* regulatory sequences without any prior DNA methylation at its coding sequence. Hence, our study shows that PTGS is dispensable for *de novo EVD* silencing. Although we cannot rule out that PTGS might facilitate the initiation of TGS, or control TE activity until then, initiation of epigenetic silencing can take place in its absence.

## Introduction

Due to their mobile nature, transposable elements (TEs) pose a threat to genome integrity and can cause mutations compromising host fitness (McClintock, 1984; Bennetzen & Wang, 2014; Schubert & Vu, 2016; Bourque *et al*, 2018). To prevent their activity, TEs are mostly transcriptionally repressed across genomes through the action of epigenetic silencing mechanisms (Lippman *et al*, 2004; Allshire & Madhani, 2018). In *Arabidopsis thaliana*, the best studied model for plants, DNA methylation (5-methylcytosine; 5mC) and histone H3 lysine-9 di-methylation (H3K9me2) cooperatively mediate transcriptional gene silencing (TGS) of TEs (Bernatavichute *et al*, 2008). Once established, 5mC and H3K9me2 patterns are propagated across generations to ensure transgenerational silencing of TEs. Maintenance of cytosine methylation depends on its sequence context, CG, CHG, or CHH (where H can be any nucleotide besides G). METHYLTRANSFERASE 1 (MET1) preserves 5mC in the CG context after each DNA replication cycle (Mathieu *et al*, 2007). Maintenance in CHG and CHH contexts occurs through a self-reinforcing loop with H3K9me2 (Chan *et al*, 2005; Du *et al*, 2012; Law *et al*, 2013; Du *et al*, 2014). In addition, DNA methylation deposition, particularly in the CHH context, can be mediated by the RNA-directed DNA methylation (RdDM) pathway. Canonical RdDM relies on the action of two plant specific RNA polymerases, PolIV and PolV. On the one hand, PolIV is recruited to TE loci through the histone mark H3K9me2. PolIV transcripts are converted into dsRNAs. These are further processed into 24-nt siRNA by DICER-LIKE 3 (DCL3), which are loaded into ARGONAUTE (AGO) 4/6-clade proteins. On the other hand, PolV is recruited to DNA-methylated TE loci. PolV provides the scaffold/target transcript for loaded AGO proteins, guiding the deposition of DNA methylation in all cytosine contexts (Law & Jacobsen, 2010; Kuhlmann & Mette, 2012; Matzke & Mosher, 2014; Erdmann & Picard, 2020). Given the dependency on PolIV-derived siRNAs, this branch of RdDM is also known as PolIV-RdDM.

While the maintenance of TE silencing relying on pre-existing heterochromatic marks is well described, the deposition of *de novo* silencing marks on active, proliferative transposable elements remains poorly understood. To gain insight into the initial molecular events by which plants recognize and silence active TEs, several studies have investigated host responses to environmentally, chemically, developmentally, or genetically induced TE reactivation (Teixeira *et al*, 2009; Slotkin *et al*, 2009; Mirouze *et al*, 2009; Reinders *et al*, 2009; Ito *et al*, 2011; Thieme *et al*, 2017). One of the best studied TEs in Arabidopsis is the *Ty1/Copia* long terminal repeat (LTR) retrotransposon *ÉVADÉ (EVD; Copia93)* (Mirouze *et al*, 2009). *EVD* is a functional, low copy TE in the reference Col-0 Arabidopsis ecotype and mainly regulated through CG methylation. Therefore, it can be released from silencing through loss of MET1 or the chromatin remodeler DECREASE IN DNA METHYLATION 1 (DDM1). Once lost, CG methylation cannot be reestablished, thus reactivated *EVD* remains active even after the reintroduction of functional (wild type) alleles of either *MET1* or *DDM1* and can rapidly increase in copy number all over the genome (Mathieu *et al*, 2007; Reinders *et al*, 2009; Mirouze *et al*, 2009). Owing to such property, *EVD* has quickly become a model system to study retrotransposon biology, TE bursts, and *de novo* silencing phenomena (Mirouze *et al*, 2009; Tsukahara *et al*, 2009; Marí-Ordóñez *et al*, 2013; Oberlin *et al*, 2017).

The *EVD* genome colonization and silencing cycle can be divided in well-defined stages. First, upon *EVD* reactivation, post-transcriptional gene silencing (PTGS) acts as initial host response (**Fig. 1A**) (Marí-Ordóñez *et al*, 2013; Oberlin *et al*, 2022). *EVD* PTGS is the result of its transcriptional and translational strategy to complete its transposition cycle. Due to an alternative splicing event, *EVD* produces two transcripts: (i) a full-length polycistronic mRNA (*fl-GAG-POL*) encoding for both its structural (Gag nucleocapsid) and catalytic (Pol) components; (ii) a short, Gag only transcript (*shGAG*), which is preferentially translated to generate the molar excess of Gag-to-Pol needed for the formation of virus-like particle (VLP) (Oberlin *et al*, 2017; 2022). A ribosome stalling event triggered during *shGAG* translation leads to the cleavage of the transcript. The resulting 3’ RNA fragment becomes a substrate for the RNA-DEPENDENT RNA POLYMERASE 6 (RDR6) to produce double-stranded (ds)RNA, further processed by DICER-LIKE 4 (DCL4) into *GAG*-derived 21-nt siRNA (Oberlin *et al*, 2022). Albeit PTGS does reduce *EVD GAG* mRNA and protein levels, it does not prevent *EVD* transposition (Marí-Ordóñez *et al*, 2013; Oberlin *et al*, 2022). Next, *EVD* continues to increase its copy number across generations. When a threshold of 40-50 *EVD* copies per genome is reached, the excess of dsRNA produced by RDR6 is eventually processed by DCL3, giving rise to a population of *GAG*-derived 24-nt siRNAs (**Fig. 1A**). Loaded into AGO4-clade proteins, they guide the deposition of DNA methylation at *EVD-GAG*-coding sequences in a non-canonical RdDM pathway also known as RDR6-RdDM (Nuthikattu *et al*, 2013; Marí-Ordóñez *et al*, 2013). Finally, following *GAG* methylation, a switch from PTGS to transcriptional gene silencing (TGS) takes place. *EVD* TGS is characterized by the installation of PolIV-RdDM and the production of 24-nt siRNAs from *EVD-LTR* sequences, mediating DNA methylation of its regulatory sequences and the *de novo* TGS of new *EVD* copies (**Fig. 1A**) (Marí-Ordóñez *et al*, 2013).

**Figure 1:**
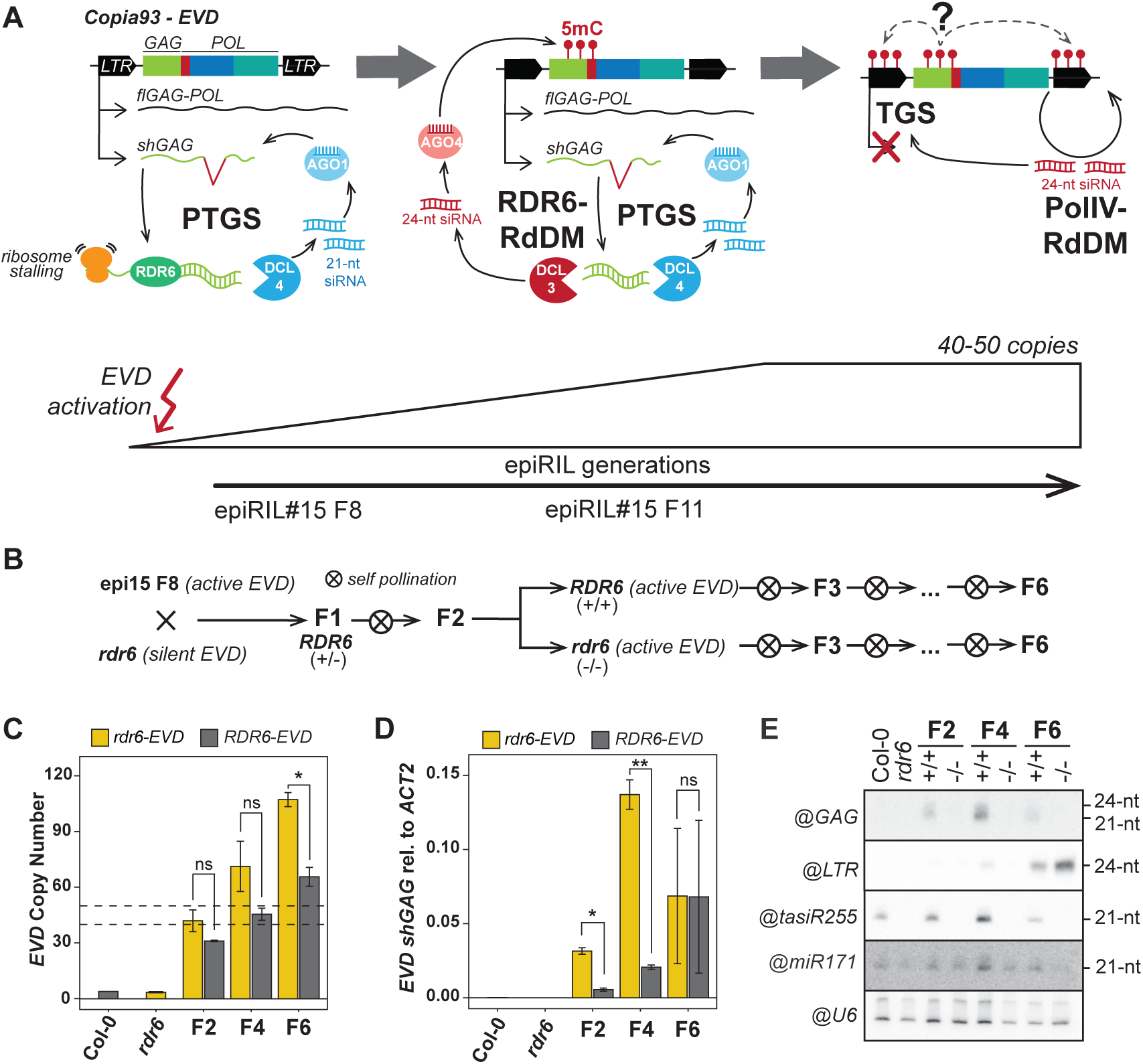
Introgression and characterization of *EVD* in the *rdr6* mutant background. **A)** Schematic representation of the three *EVD* silencing steps. Upon *EVD* reactivation, ribosome stalling during translation of *EVD shGAG* transcript triggers PTGS. 21-nt siRNA produced through RDR6 and DCL4 and loaded into AGO1. With increasing *EVD* copies across generations, the excess of dsRNA produced by RDR6 is processed by DCL3 to generate 24-nt siRNAs. Loaded into AGO4, *shGAG* siRNAs trigger DNA methylation (5mC) through RDR6-RdDM at GAG coding sequences without silencing. At 40-50 copies per genome, TGS is installed through Pol IV-RdDM, coincidentally with the appeareance of DNA methylation and 24-nt siRNAs on the LTR sequences. **B)** Crossing scheme to generate *rdr6* mutant lines with active *EVD.* F2 plants were genotyped to select homozygous WT and mutant RDR6 lines, propagated through selfing until the F6 generation. **C)** *EVD* copy number analysis by qPCR in RDR6-*EVD* and *rdr6-*EVD lines at generations F2, F4 and F6, using the *EVD*-GAG sequence as target. **D)** qPCR analysis of *shGAG* expression normalized to *ACT2* in *EVD*-RDR6 and *EVD*-*rdr6* lines at generations 2, 4 and 6. Two biological replicates are represented, error bars show standard error in C,D (n.s p>0.05, *p<0.05, **p<0.01, ***p<0.005, two-sided t-test between indicated samples). **E)** RNA blot analysis of *EVD* siRNAs against GAG and LTR in RDR6 and *rdr6* lines with active *EVD* at generations F2, F4 and F6. tasiR255 probe is used as control for RDR6 mutation, miR171 and snoRNA U6 are shown as loading controls. WT Col-0 and *rdr6* with no reactivated *EVD* are shown as negative control for *EVD* activity.

These observations have led to a model under which RDR6-RdDM is thought to contribute to the switch from PTGS to TGS by initiating the deposition of DNA methylation on *EVD GAG*-coding sequences. PTGS-derived siRNAs have been shown to play a role in restoring DNA methylation patterns in DNA methylation mutants (Teixeira *et al*, 2009; Nuthikattu *et al*, 2013; McCue *et al*, 2015) or during key developmental processes involving epigenetic reprogramming (Slotkin *et al*, 2009). Hence, RDR6-RdDM has been suggested as essential step to gap the transition from PTGS to TGS in the process of *de novo* transposon silencing (Marí-Ordóñez *et al*, 2013; Nuthikattu *et al*, 2013; McCue *et al*, 2015; Panda *et al*, 2016). More recently, using an *EVD* transgenic system, a mechanism for such transition has been proposed, under which AGO4, loaded with RDR6-derived siRNAs, interacts with PolII *EVD* transcripts at *EVD* loci to recruit PolV and downstream silencing components to ignite self-sustained PolIV-RdDM (Sigman *et al*, 2021). However, the requirement of PTGS for the initiation of TGS during a TE colonization event has never been experimentally tested. Furthermore, upon genome-wide loss of DNA methylation, the majority of reactivated TEs triggering RDR6-dependent PTGS are decayed TE-remnants incapable of transposition, while those intact enough to engage in translation do not (Oberlin *et al*, 2022). This has casted doubt on the universality of PTGS as a sensor of active TEs and initiator of epigenetic silencing during a TE colonization event.

In this study we address the necessity of PTGS and RDR6-RdDM in the process of *de novo* silencing of transposable elements. To achieve this, active *EVD* was introduced into *RDR6* mutant plants to prevent RDR6-RdDM. *EVD* activity and silencing status were monitored over several generations in wild-type (WT) and *rdr6* lines. While the mutation of *RDR6* successfully prevented the production of siRNAs involved in RDR6-RdDM, silencing of *EVD* through installation of PolIV-RdDM in its LTRs was achieved in both, WT and mutant background. Therefore, PTGS is not essential for the installation of epigenetic silencing at active *EVD* copies. Given that *EVD* presents a unique case in triggering PTGS upon reactivation in the first place, we suggest that PTGS and RDR6-derived siRNA play a role in limiting *EVD* expression and transposition rather than initiating *de novo* silencing.

## Results

### Absence of RDR6 does not prevent the production of *EVD-LTR* 24-nt siRNAs

To investigate the role of PTGS and RDR6-RdDM in the initiation of canonical RdDM on *EVD*, and given that RDR6 is responsible for the production of *EVD* siRNA during the PTGS phase (Oberlin *et al*, 2017; 2022), , *rdr6-15* mutant plants were crossed to the 8^th^ generation of the *met1*-derived epigenetic recombinant inbred line (epiRILs) number 15 (epi15 F8), a generation in which RDR6-RdDM has not yet been activated (**Fig. 1B**), (Marí-Ordóñez *et al*, 2013; Oberlin *et al*, 2022). This minimized the probability of introducing active *EVD* copies with *GAG* DNA methylation in the cross to *rdr6*. In the second generation (F2), two homozygous RDR6 wild-type (RDR6) and two mutant (*rdr6*) plants were selected by genotyping (hereafter referred to as RDR6-*EVD* and *rdr6-EVD,* respectively). They were allowed to self-pollinate in order to bulk-propagate two independent WT and two independent muntant populations until the 6^th^ generation (F6), allowing *EVD* to colonize the genome (**Fig. 1B**). The silencing status of *EVD* in the two independent respective populations was monitored in bulks of 8 plants at generations F2, F4, and F6 by assessing the *EVD* copy number and expression levels through qPCR as well as the siRNA profile by RNA blot.

*EVD* copy numbers consistently increased across generations, confirming the inheritance of active, transposition-competent *EVD* copies from epi15 (**Fig. 1C**). *EVD* copies accumulated at a higher rate in *rdr6*-*EVD* background than in *RDR6*-*EVD*. The estimated copy number was consistently higher in *rdr6-EVD* than in *RDR6*-*EVD* plants, and the difference was significant in the F6, where loss of RDR6 activity allowed the accumulation of over 100 *EVD* copies (**Fig. 1C**), the absence of RDR6 activity allowed the accumulation of over 100 *EVD* copies in the F6 generation (**Fig. 1C**). Consistent with an increasing copy number, *EVD* transcripts were also more abundant up to the 4^th^ generation, particularly in *rdr6-EVD*. However, at the F6, large variations between biological replicates were observed in both *RDR6-EVD* and *rdr6-EVD* (**Fig. 1D**). This variation was observed consistently for the expression of both *shGAG* and *flGAG-POL* isoforms (**Fig. 1D**; **Supp. Fig. 1**), suggesting a biological rather than a technical origin. The decrease and broad variation in *EVD* expression compared to the F4 (and despite the increase in copy number) suggested that TGS had started to take place. This was expected for the *RDR6*-*EVD* as the bulk of F6 individuals has already exceeded the 40-50 copy number limit, previously demonstrated to lead to TGS (Marí-Ordóñez *et al*, 2013). However, that result was unanticipated and intriguing for the *rdr6-EVD* lines and initiated the analysis of the potential *EVD* silencing mechanism by investigating the small RNA profile by RNA blots.

Although the transition of *EVD* from PTGS to TGS has been demonstrated to take place at the individual plant level, the detection of *EVD-GAG* and *EVD-LTR* siRNA in bulked material can also be used as proxy for the assessment of *EVD* silencing stage at each generation (Marí-Ordóñez *et al*, 2013). As expected from the dependency of *GAG*-derived siRNA on *EVD* expression, *EVD-GAG* siRNAs were detected in RDR6-*EVD* throughout generations, mirroring *EVD* expression levels (**Fig. 1E**). In these plants, 24-nt *EVD-LTR* siRNAs, involved in PolIV-RdDM and transcriptional silencing of *EVD,* were first detected in the F4 and increased in the F6 generation, coinciding with the decrease of *EVD* expression and *GAG* siRNAs (**Fig. 1E**). In contrast, as expected in the absence of RDR6, no RDR6-derived *EVD-GAG* siRNAs and trans-acting (ta)siRNAs were detectable in *rdr6-EVD* plants, *EVD-LTR* siRNAs were present in the F6 generation (**Fig. 1E**). Thus, the lack of PTGS and RDR6-derived *EVD-GAG* siRNA did not prevent the appearance of *EVD-LTR* 24-nt siRNAs, previously associated with *EVD* TGS.

### RDR6-dependent *EVD-GAG* siRNAs are dispensable for *EVD* TGS

The presence of 24-nt *EVD-LTR* siRNAs, hallmark of successful PolIV-RdDM installation, indicated that silencing of *EVD* copies through TGS was likely taking place in both RDR6-*EVD* and *rdr6-EVD* backgrounds. Previous work had shown that *EVD-GAG* siRNAs (PTGS) and *EVD-LTR* 24-nt siRNAs (PolIV-RdDM) are mutually exclusive at the individual level. While both can be detected in bulks of plants, individuals with active or silenced *EVD* produce *GAG* or *LTR* siRNAs respectively as the transition to TGS has been shown to impact most if not all copies within one generation at the individual level (Marí-Ordóñez *et al*, 2013). The detection of both in RDR6-*EVD* bulks suggested that a subset of the individual plants successfully silenced *EVD* through TGS, pointing to a different silencing status in individual F6 plants within the bulked samples of both lines.

To investigate *EVD* silencing at the level of individual plants, and whether TGS had taken place in the absence of RDR6, ten *RDR6*-*EVD* and ten *rdr6-EVD* individuals from the F6 generations plants were selected. In *RDR6*-*EVD*, 24-nt *EVD-LTR* siRNAs were detected in all but two individuals, #2 and #9, where only *EVD-GAG* siRNAs were present (**Fig. 2A**). The presence of 24-nt *EVD-LTR* siRNAs was associated with a lower expression of *EVD* (**Fig. 2B**). Similarly, 24-nt *EVD-LTR* siRNAs were also detected in 6 out of 10 *rdr6-EVD* individuals (**Fig. 2A**) and the presence of 24-nt siRNAs correlated with a corresponding loss of *EVD* expression (**Fig. 2B**). Hence, silencing of *EVD* and production of associated *LTR* 24-nt siRNAs can take place in absence of RDR6-dependent *GAG* siRNAs.

**Figure 2:**
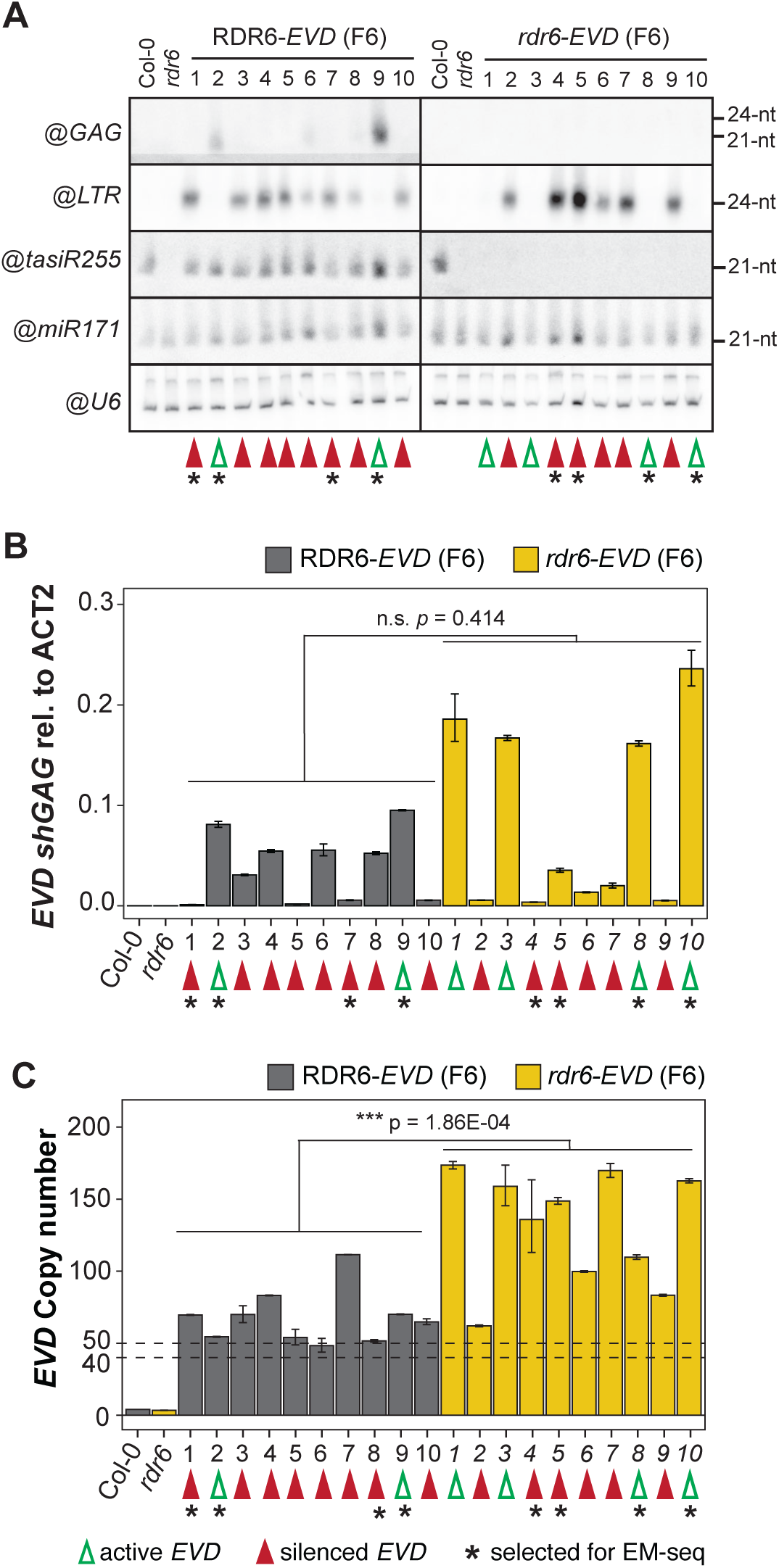
Characterization of *EVD* silencing status in *RDR6-* and *rdr6-EVD* F6 individuals. **A)** RNA blot analysis of *EVD* siRNAs against GAG and LTR in 10 F6 individual plants of RDR6-*EVD* and *rdr6-EVD* lines. tasiR255 probe is used as control for RDR6 mutation, miR171 and snoRNA U6 are shown as loading controls. WT Col-0 and *rdr6* with no reactivated *EVD* are shown as negative controls for *EVD* activity and expression. **B)** Analysis of *EVD shGAG* expression of the same individuals investigated in A by qPCR, normalized to *ACT2.* **C)** *EVD* copy number analysis by qPCR of the same individuals investigated in A and B, using the *EVD*-GAG sequence as target. Three technical replicates are represented, error bars show standard error in B and C (n.s p>0.05, *p<0.05, **p<0.01, ***p<0.005, two-sided t-test between indicated samples). Green and red arrows indicate individuals with active and silenced *EVD* copies respectively, selected individuals for subsequent EM-sequencing are indicated by an asterisk in A, B, C.

### *EVD* silencing in the absence of RDR6 does not correlate with copy number

*EVD* switch to TGS has been experimentally established at around 40-50 copies in both *met1* and *ddm1* epiRILs, coinciding with the upper limit of natural variation for *COPIA93* copies found within *Arabidopsis* ecotypes (Marí-Ordóñez *et al*, 2013; Quadrana *et al*, 2016). To assess whether a similar threshold applied in the absence of RDR6, we quantified *EVD* copy number in the same *RDR6-EVD* and *rdr6-EVD* F6 individuals used for *EVD* siRNAs and expression.

In agreement with the copy number threshold above which *EVD* TGS takes place, most *RDR6-EVD* individuals displayed homogenous copy number only slightly above this range. In *rdr6-EVD* plants, however, *EVD* silencing had taken place at a more variable copy number (**Fig. 2C**). Many plants displayed higher copy number, as previously observed in the bulk analysis (**Fig. 1C**); some individuals had switched to TGS at copy numbers just above the threshold (*rdr6-EVD* #2), while others did so at a copy numbers well above 100 (*rdr6-EVD* #4, 5, 7) (**Fig. 2C**). Nonetheless, *EVD* remained active in other individuals with copy numbers above 150 (*rdr6-EVD* #1, 3, 10) (**Fig. 2C**). Consequently, in the absence of RDR6 activity, no clear copy number threshold for TGS installation was observed. Thus, in the *rdr6* background, silencing of active *EVD* might be stochastic once the 40-50 copy number threshold is exceeded or depend on other factors not considered so far.

### DNA methylation of *EVD-GAG* is dispensable for the transition to TGS

The copy number threshold is likely determined by the point at which the cumulative *EVD* expression from all new insertions causes DCL3 to process *shGAG* RDR6-derived dsRNA to initiate RDR6-RdDM (Marí-Ordóñez *et al*, 2013). Although *EVD* switch to TGS took place in absence of RDR6 and associated *GAG* siRNAs, we could not rule out the processing of *shGAG* transcripts by one of the other Arabidopsis RDR proteins, producing siRNA levels undetectable by Northern blots, but sufficient to induce the *EVD-GAG* DNA methylation believed to initiate PolIV-RdDM.

To corroborate the installation of PolIV-RdDM at *EVD LTRs* and address whether DNA methylation was independently of RDR6-generated siRNAs still deposited at *EVD-GAG*, or another coding region, *EVD* DNA methylation was assessed by whole-genome Enzymatic Methyl-sequencing (EM-seq) (Feng *et al*, 2020) on F6 individuals before and after the transition to TGS. We selected lines with active *EVD* (two *RDR6*-*EVD* and two *rdr6-EVD* individuals with high *EVD* expression and no 24-nt *EVD-LTR* siRNAs), as well as lines with silenced *EVD* (two RDR6-*EVD* and two *rdr6-EVD* individuals producing 24-nt *EVD-LTR* siRNAs and low *EVD* expression) (**Fig. 2A, B**; marked by asterisks). EM-seq on wild-type (*Col-0*) and *rdr6* plants was performed as control for endogenous *EVD* methylation in the respective background. Overall, a 30-50X genome coverage with conversion rates above 99.8% was obtained for all samples (**Supplementary Fig. S2**). Because short read sequencing technology makes it challenging to estimate methylation levels along individual *EVD* insertions (∼5 kb), EM-seq reads were mapped to a single fictitious *EVD* locus to estimate average methylation levels.

In *RDR6-EVD* with active *EVD,* low levels of *GAG* methylation in all cytosine contexts were observed. In contrast, the equivalent *rdr6-EVD* lines displayed near absence of DNA methylation (**Fig. 3A, B**). Furthermore, DNA methylation increased upon *EVD* silencing in *RDR6-EVD* but remained low after the *EVD* silencing in *rdr6-EVD* (**Fig. 3A**), indicating that absence of RDR6 was sufficient to abolish *EVD-GAG* DNA methylation, not only during *EVD* proliferation, but also after silencing. In addition, DNA methylation levels in *rdr6-EVD* lines remained low across other *EVD* coding regions in *rdr6-EVD* lines with active or silenced *EVD,* compared to *RDR6-EVD* (**Supplementary Fig. S3A, B**), ruling out that the switch to TGS is triggered through non-canonical-RdDM activity at other regions in the absence of RDR6.

**Figure 3:**
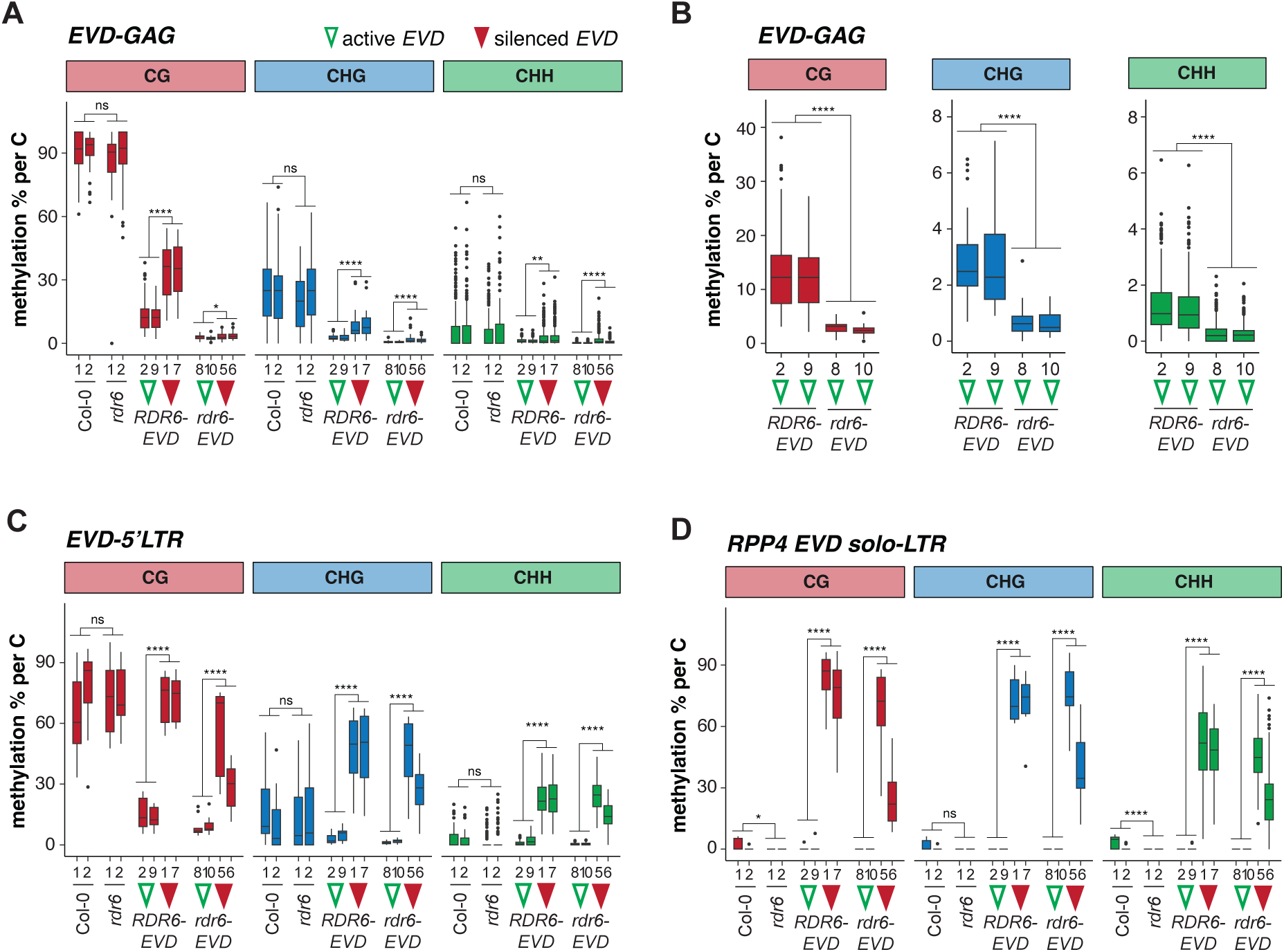
DNA methylation of active and silenced *EVD* in *RDR6-* and *rdr6-EVD* lines. EM-seq analysis of DNA methylation (as methylation % per cytosine) in CG, CHG and CHH contexts in WT Col-0, *rdr6* and in *RDR6-* and *rdr6-EVD* F6 individuals with active and silenced *EVD* (marked with empty green and filled red arrowheads respectivey, nubers indicate same individuals as in Figure 2) in: **A**, *EVD-GAG*; **B**, *EVD-GAG* but only in *RDR6-* and *rdr6-EVD* F6 individuals with active *EVD*; **C**, *EVD-5’LTR* and; **D**, *EVD solo-LTR* in *RPP4* (AT4G16869) promoter. In all boxplots: median is indicated by a solid bar, the boxes extend from the first to the third quartile and whiskers reach to the furtherst values withinn 1.5 times the interquartile range. Dots indicate outliers, as data points outside of the above range. Wilcoxon rank sum test adjusted p-values between indicated groups of samples: *: p<0.05, **: p<0.01, ***: p<0.001, ****: p<0.0001, ns: non-significant (p≥0.05) Wilcoxon rank sum test between indicated groups of samples.

We next investigated DNA methylation at *EVD-LTRs* to confirm that presence of *LTR* 24-nt siRNA were *bona fide* indicators of PolIV-RdDM and the switch to *EVD* TGS. In both *RDR6-EVD* and *rdr6-EVD* lines with active *EVD*, DNA methylation at the *LTRs* was very low in all three contexts, while methylation levels were increased in those with silenced *EVD* (**Fig. 3C**). Furthermore, methylation levels at CHG and specifically at CHH were higher than those of the parental *EVD* copy in the wild-type *Col-0* and

*rdr6* controls (**Fig. 3C, Supplementary Fig. S3C**), confirming the successful installation of PolIV-RdDM following the *EVD* burst, in contrast to RdDM-independent DNA methylation maintenance of *EVD* prior to its reactivation. To further validate that the switch to TGS through PolIV-RdDM installation had taken place, we inspected the methylation status of the *EVD*-derived solo-*LTR* present in the promoter of *RECOGNITION OF PERONOSPORA PARASITICA 4* (*RPP4*, *AT4G16860*), which can be methylated *in trans* by *EVD-LTR* 24-nt siRNAs following an *EVD de novo* silencing event (Marí-Ordóñez *et al*, 2013). Indeed, *EVD solo-LTR* was only methylated in the *RDR6-EVD* and *rdr6-EVD* lines where *EVD* had been silenced (**Fig. 3D**).

Hence, the switch to TGS through PolIV-RdDM does neither require RDR6-RdDM nor the deposition of DNA methylation on *EVD-GAG* or other coding regions, to successfully silence *EVD*.

### EM-seq paired-end sequencing data allows mapping of new *EVD* insertions

Once PolIV-RdDM is established, *EVD* silencing and associated deposition of DNA methylation impacts new copies genome-wide through the trans-activity of *EVD LTR* 24-nt siRNAs (Marí-Ordóñez *et al*, 2013). However, in *EVD-*silenced *rdr6-EVD* lines, *LTR* methylation levels were lower in the CG context for line #5 and in all contexts for line #6 than those in *RDR6-EVD* (**Fig. 3C, Supplementary Fig. S3C**). To investigate the homogeneity of DNA methylation at individual *EVD* insertions, and to ask whether it is influenced by the absence of RDR6-RdDM, we took advantage of discordant paired-read mates in our EM-seq data, where one of the read mates mapped to *EVD LTRs* and the other elsewhere in the genome, to identify and locate new *EVD* insertions (Gilly *et al*, 2014; Stuart *et al*, 2016; Quadrana *et al*, 2016; 2019) (**Fig. 4A**, **Supplementary Fig. S4**) and to assess their methylation levels. Only new *EVD* insertions supported by three or more discordant paired-read mates from both of *EVD LTRs* were considered. Simultaneously, concordant paired-read mates mapping to *EVD* were used to independently estimate *EVD* copy numbers through the increase in *EVD* sequencing coverage due to additional insertions in *RDR6-* and *rdr6-EVD* relative to the Col-0 reference genome (Yoon *et al*, 2009; Quadrana *et al*, 2016) (**Fig. 4A**, **Supplementary Fig. S4**).

**Figure 4:**
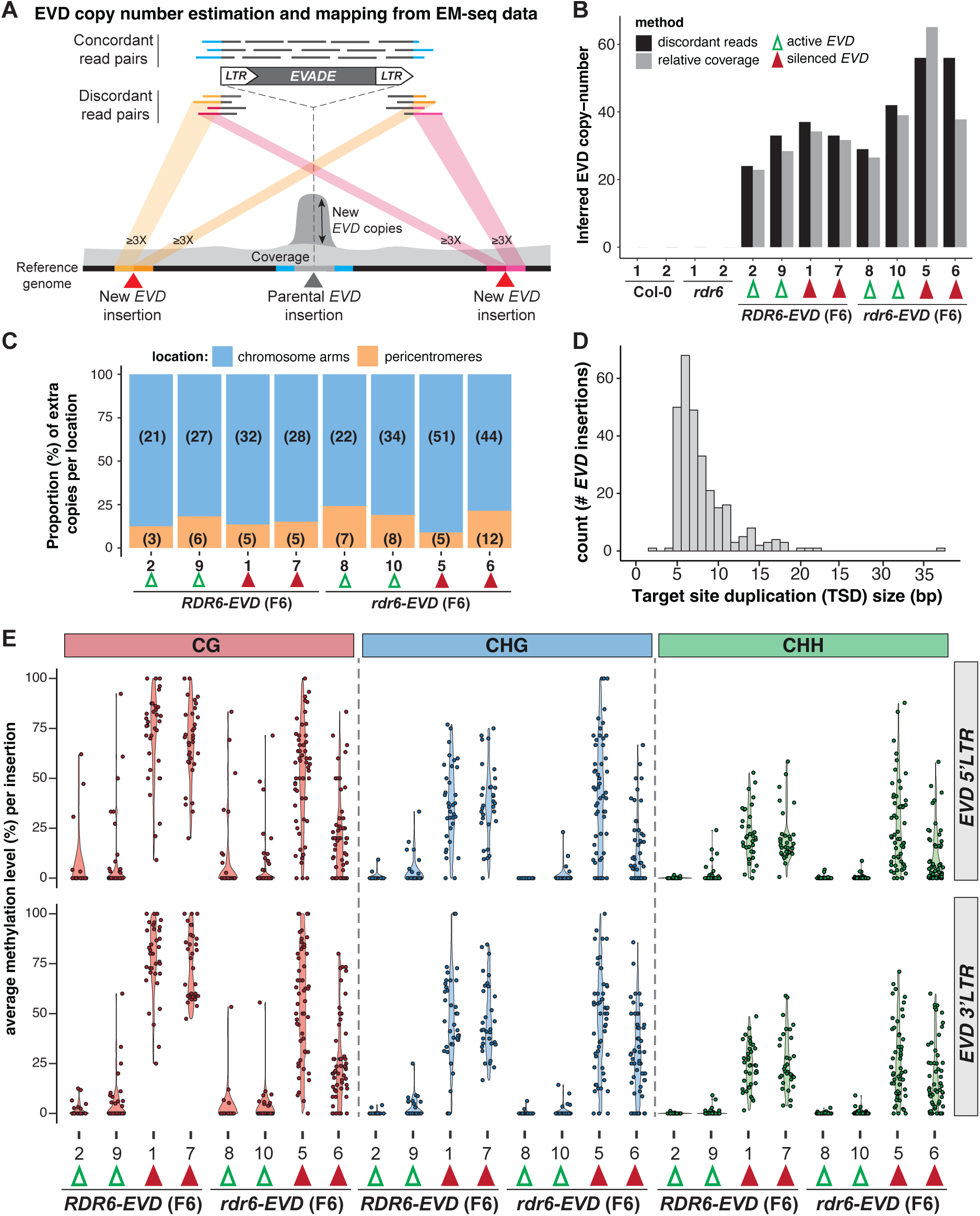
*LTR* DNA methylation of individual new *EVD* inertions in *RDR6-* and *rdr6-EVD* lines. **A)** Schematic representation of the strategy used to quantify and map new *EVD* insertions from EM-seq data. *EVD* copy number was estimated using: i) the increased *EVD* coverage of concordant paired read mates in EM-seq data, consequence of *EVD* transposition or, ii) mapping new *EVD* inserions through discordant read pairs mapping to *EVD* and elswhere in the genome. New insertions had to be supported by at least three discordant read pairs from each border to be considered. **B)** Inferred *EVD* copy number using either relative coverage or discordant reads in WT Col-0, *rdr6* and in *RDR6-* and *rdr6-EVD* F6 individuals with active and silenced *EVD* (marked with empty green and filled red arrowheads respectivey, numbers indicate same individuals as in Figures 2 and 3). **C)** Number and relative proportion (in %) of mapped new *EVD* insertions in pericentromeric or chromosomic arm locations in each indicated sample. Numbers of new *EVD*insertions at each location are indicated in brackets. **D)** Histogram of the size (in bp) distribution of target site duplications at all new *EVD* insertions. **E)** *5’* and *3’LTR* average DNA methylation levels in each cytosine context for individual new *EVD* insertions in *RDR6-* and *rdr6-EVD* F6 individuals with active and silenced *EVD*.

While no new *EVD* insertions were obtained in Col-0 and *rdr6* controls, estimation methods yielded consistent increased *EVD* copy numbers in lines where *EVD* had proliferated (**Fig. 4B**). As shown above (**Fig. 2C**), more *EVD* insertions were found in *rdr6-EVD* than in *RDR6-EVD* (**Fig. 4B**). In all lines, 75% or more of new insertions were mapped to chromosome arms (**Fig. 4C**, **Supplementary Fig. S5**), as expected from the integration preference of *EVD* into gene-rich regions (Quadrana *et al*, 2019). Furthermore, most new *EVD* insertions caused short sequence duplications, known as target site duplications (TSD), of 5-8 bp (**Fig. 4D**). This falls within the TSD range expected for LTR-RTE in plants , specifically a 5-bp TSDs as obtained here for *EVD* (Quadrana *et al*, 2016; Orozco-Arias *et al*, 2019; Jedlicka *et al*, 2019; Roquis *et al*, 2021). Therefore, the new *EVD* insertions mapped from EM-seq data displayed features of *bona fide* new *EVD* transposition events.

We noticed that *EVD* copy numbers quantified from EM-seq data were lower than the quantification by qPCR. This discrepancy might originate from inaccuracy of qPCR quantifications, including potential amplification of *EVD* extra-chromosomal cDNA, or from limited discordant read-mates coverage in the EM-seq data. Nonetheless, defined insertions identified in all samples allowed to investigate methylation levels at individual *LTRs* of new *EVD* insertions.

### DNA methylation is not homogeneously deposited across new *EVD* insertions

As observed in our global analysis of *EVD* DNA methylation (**Fig. 3**), both *LTRs* of individual new *EVD* insertions gained DNA methylation in all contexts in *EVD-*silenced lines compared to those with active *EVD*. However, the *LTR* DNA methylation levels of individual *EVD* insertions were less homogeneous than expected, displaying a broad range in all contexts. Notwithstanding, we observed that the levels of DNA methylation in the three contexts were more heterogenous in *rdr6-EVD* lines than in the equivalent *RDR6-EVD* ones (**Fig. 4E**). This was most remarkable in the CG context, where most insertions in *RDR6-EVD* lines with silenced *EVD* displayed CG methylation levels above 50%, whereas several insertions in *rdr6-EVD* display lower or no methylation, especially in the *rdr6-EVD* line #6. A similar trend was observed in the CHG and CHH contexts, where some insertions displayed lower or no methylation in the *rdr6-EVD* lines after the switch to PolIV-RdDM (**Fig. 4E**). This could be a consequence of the absence of potential priming for the switch to TGS provided by *EVD-GAG* DNA methylation but might also reflect a difference in the timing of the switch. *EVD-LTR* 24-nt siRNAs were already detected at the 4^th^ generation in *RDR6-EVD* but not in *rdr6-EVD* (**Fig. 1E**). Thus, in the later, *EVD* likely had been under PolIV-RdDM for less generations. This might be the case especially in the *rdr6-EVD* #6 line, where *RPP4 solo_LTR* methylation levels are also lower than in the other *EVD-* silenced lines (**Fig. 3D**). Nonetheless, the gain of DNA methylation at most new *EVD* insertions further supports that PolIV-RdDM gets installed at *EVD-LTRs* in the absence of RDR6-RdDM.

### RdR6-RdDM is insufficient for *EVD* silencing in the absence of PolIV-RdDM

The above results indicated that, independently of RDR6-RdDM and *EVD-GAG* methylation, *EVD* silencing took place through the initiation of PolIV-RdDM at *EVD-LTR*s. Nonetheless, given that PolIV-RdDM requires pre-existing epigenetic marks for its recruitment, *EVD-LTR* 24-nt siRNAs might be a consequence of a preceding *EVD* silencing event rather than the cause. While both PolIV and PolV are essential for PolIV-RdDM, only PolV is required for RDR6-RdDM, if siRNAs are provided through PTGS (Nuthikattu *et al*, 2013; McCue *et al*, 2015; Taochy *et al*, 2019; Sigman *et al*, 2021). Hence, to test the dependency of *EVD* TGS in PolIV-RdDM and, at the same time, explore if RDR6-RdDM could lead to TGS independently of PolIV-RdDM, we introduced active *EVD* in the mutants *nrpd1 (*PolIV largest subunit mutant) and *nrpe1* (PolV largest subunit mutant), following the same strategy as for *rdr6*. This time, however, to ensure the presence of *EVD-GAG* methylation before the loss of RdDM, an epi15 F11 generation plant, already undergoing RDR6-RdDM (Marí-Ordóñez *et al*, 2013) (**Fig. 1A**), was used as *EVD* donor. Again, wild-type and mutant plants were selected in the F2 and propagated to F6 (**Supp. Fig. 6A**).

While lines carrying wild-type or mutant alleles were indistinguishable with respect to *EVD* copy number or expression in the F2 (**Fig. 5A, B**), *EVD* reached higher copy numbers, surpassing the 40-50 copy number threshold, in the following generations of both mutants (*nrpd1-EVD* and *nrpe1-EVD*) than in the lines carrying the corresponding wild-type alleles (*NRPD1-EVD* and *NRPE1-EVD*) (**Fig. 5A**). Similarly to the results obtained with *RDR6-EVD* lines, once the threshold was reached in lines with wild-type alleles, *EVD* expression was reduced. However, in both mutant backgrounds, *EVD* remained transcriptionally active (**Fig. 5B**), indicating that TGS was not installed in the absence of PolIV-RdDM. Furthermore, investigation of *EVD* small RNA patterns in three independent bulks of plants at the F6 generation confirmed the presence of *EVD-LTR* 24-nt siRNAs in both *NRPD1-* and *NRPE1-EVD* lines. On the contrary, in *nrpd1-* and *nrpe1-EVD* lines, only *EVD-GAG* siRNAs were detected (**Fig. 5C, D**), consistent with *EVD* expression triggering PTGS.

**Figure 5:**
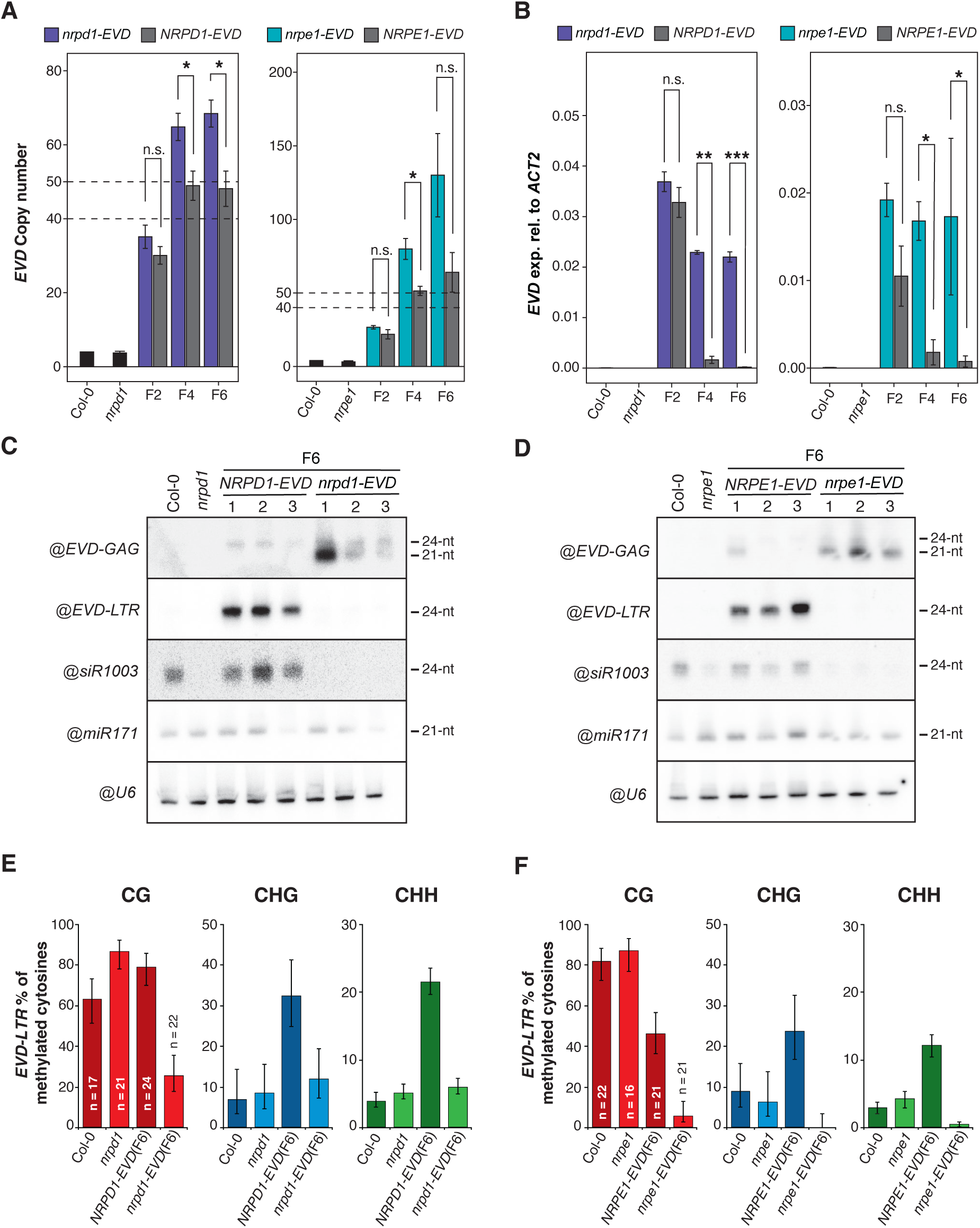
Characterization of *EVD* proliferation and silencing in RdDM mutants. **A)** *EVD* copy number analysis by qPCR in *NRPD1-EVD* and *NRPE1-EVD* lines, in both WT (grey) and mutant (colored) backgrounds, at generations F2, F4 and F6, using the *EVD*-GAG sequence as target. **B)** qPCR analysis of *shGAG* expression normalized to *ACT2* in *NRPD1-EVD* and *NRPE1-EVD* lines, in both WT (grey) and mutant (colored) backgrounds, at generations F2, F4 and F6. Three biological replicates are represented, error bars show standard error in A, B (n.s p>0.05, *p<0.05, **p<0.01, ***p<0.005, two-sided t-test between indicated samples). **C – D)** RNA blot analysis of *EVD* siRNAs against GAG and LTR in the F6 generation of 3 independent WT and mutant lines of *NRPD1-EVD* (**C**) and *NRPE1-EVD* (**D**). WT Col-0 and *nrpd1* or *nrpe1* with no reactivated *EVD* are shown as negative control for *EVD* activity. siR1003 probe is used as control for *NRPD1* and *NRPE1* mutations, miR171 and snoRNA U6 are shown as loading controls. **E – F)** Bisulfite-PCR DNA methylation analysis at *EVD-*LTR sequences in the F6 generation of *NRPD1-EVD* (E) and *NRPE1-EVD* (F) lines, in both WT (darker shade) and mutant (lighter shade) backgrounds. Col-0, *nrp1d* and *nrpe1* were used as controls. n: number of clones analyzed. Error bars represent 95% confidence Wilson score intervals.

Methylation of *EVD* insertions was assessed in F6 bulks by Sanger sequencing of PCR amplicons from bisulfite-treated DNA (BS-PCR). Regarding *EVD-GAG*, CG methylation was high in both wild type and mutant lines, as expected according to MET1 maintenance and previous exposure to RDR6-RdDM (**Supp. Fig. 6B-E**). CHG and CHH methylation was present in both WT and mutant lines, albeit higher in *NRPD1-* and *NRPE1-EVD* lines than in their mutant equivalents (**Supp. Fig. 6B-E**), probably due to the increase in methylation previously observed after the installation of TGS (**Fig. 3A**). Both *nrpd1-* and *nrpe1-EVD* lines displayed similar CHG methylation levels likely inherited from the epi15 F11 and maintained independently of siRNAs. However, CHH methylation was higher in *nrpd1-EVD* than in *nrpe1-EVD* lines (**Supp. Fig. 6B-E**), in agreement with the absence of RDR6-RdDM in *nrpe1* but not in *nrpd1* mutants. Therefore, RdD6-RdDM seems to still be depositing CHH methylation in the *nrpd1-EVD* lines.

We next investigated the methylation status of *EVD-LTR*. As anticipated from the different epigenetic regulation of *EVD* before and after its mobilization and silencing, the presence of 24-nt siRNAs in *NRPD-* and *NRPE-EVD* lines correlated with increased DNA methylation levels, where CHG and CHH methylation levels surpassed those found in WT and mutant controls (**Fig. 5E, F**, **Supp. Fig. 7A, B**). Furthermore, no loss of *EVD* DNA methylation, relative to Col-0, was observed neither in *nrpd1* nor *nrpe1* control samples, confirming the absence of PolIV-RdDM regulation at the parental *EVD* insertion (**Fig. 5E, F**). In contrast, in *nrpd1-* and *nrpe1-EVD* lines, DNA methylation in all contexts remained low in conformity with the absence of *EVD-LTR* 24-nt siRNAs and TGS (**Fig. 5E, F**). However, we noticed that DNA methylation in all contexts was higher in *nrpd1-EVD* than in *nrpe1-EVD*, suggesting that continuous RDR6-RdDM activity might cause weak DNA methylation in *EVD-LTR*. Nonetheless, in the absence of PolIV-RdDM, *EVD* remained transcriptionally active up to the F6 generation despite the continuous action of PTGS and, in the case of *nrpd1-EVD*, RDR6-RdDM. Hence, PolIV-RdDM is required for *de novo* TGS initiation independently of RDR6-RdDM.

## Discussion

### PTGS is required for *GAG* methylation but dispensable for the TGS of *EVD*

Post transcriptional gene silencing mediated by RDR6-dependent siRNAs has been hypothesized to be the initiating step of TE epigenetic silencing, in particular *EVD* (Teixeira *et al*, 2009; Marí-Ordóñez *et al*, 2013; Nuthikattu *et al*, 2013; McCue *et al*, 2015; Panda *et al*, 2016; Sigman *et al*, 2021). In this study, by investigating the silencing fate of active *EVD* in wild type and mutant *RDR6* backgrounds, we shown that, although DNA methylation deposited in the *EVD-GAG* sequence is a consequence of PTGS through RDR6-RdDM and might contribute to its final silencing, in absence of RDR6 and the associated *EVD-GAG* siRNAs, transcriptional gene silencing is still achieved through the installation of PolIV-RdDM without any prior DNA methylation at *EVD* coding sequences. Therefore, PTGS and gene body DNA methylation are dispensable for de novo *EVD* silencing.

Our results add further evidence supporting that PTGS can direct the deposition of DNA methylation (Wu *et al*, 2012; Nuthikattu *et al*, 2013; McCue *et al*, 2015; Taochy *et al*, 2019). However, RDR6-RdDM is neither necessary nor sufficient to for TGS inititation, which requires PolIV-RdDM at regulatory sequences. Previous work has shown that an immobile *EVD* overexpression transgenic system under the control of the CaMV 35S promoter (*35S:EVD*), despite triggering the same PTGS response as the endogenous *EVD*, does not transition to TGS in WT plants (Marí-Ordóñez *et al*, 2013; Oberlin *et al*, 2022). This is in line with the observation that in endogenous loci and transgenes triggering strong PTGS and undergoing RDR6-RdDM, DNA methylation is unable to spread from transcribed regions to regulatory sequences to set TGS (Taochy *et al*, 2019).

### PTGS might facilitate the installation of *EVD* TGS

Nonetheless, RDR6-RdDM likely sensitizes *EVD* loci to facilitate TGS initiation, explaining the homogeneity of *EVD* copy number at which RdDM is installed at *EVD* compared to the stochasticity observed in *rdr6* mutants. Introduction of either active *EVD* or *35S:EVD* in *dcl2 dcl4* double mutants, results in the switch to TGS at 20-30 *EVD* copies and silencing of *35S:EVD* coupled with the production of 24-nt siRNAs from their promoters (Marí-Ordóñez *et al*, 2013). Hence, in the case of *EVD*, enhancing RDR6-RdDM by promoting DCL3 activity upon RDR6 dsRNA products in the absence of DCL2 and DCL4, expedites *EVD* switch from PTGS to TGS. Here, despite the lack of PolIV-RdDM, a low level of DNA methylation at *EVD-LTR* was found in RDR6-RdDM competent *nrpd1-EVD* plants. Given the close proximity of *GAG* to *5’* regulatory sequences, strong or prolonged *GAG* RDR6-RdDM might result in enough methylation adjacent to the promoter to recruit PolIV-RdDM as previously suggested (Marí-Ordóñez *et al*, 2013; Sigman *et al*, 2021). This situation may be favored for endogenous *EVD*. Although both endogenous *EVD* and *35S:EVD* produce high RDR6-dependent siRNA levels, in contrast to the ubiquitously expressed *35S* promoter, *EVD-LTR* drives expression in only a few cells (Marí-Ordóñez *et al*, 2013), where intracellular siRNAs levels might be high enough to promote DNA methylation spreading into the nearby *LTR*.

### PTGS potentially acts as an *EVD* copy number control system

Alternatively, although not mutually exclusive with the priming role of RDR6-RdDM in the switch to TGS, we propose that PTGS triggered during translation can act as a mechanism to regulate *EVD* proliferation until the switch to TGS takes place. Mutations in *RDR6* lead to increased *EVD shGAG* mRNA, protein and VLP levels, resulting in increased transposition (this study and (Lee *et al*, 2021; Oberlin *et al*, 2022)). Although the mechanisms triggering *shGAG* ribosome stalling and cleavage has not yet been elucidated, given that PTGS is not commonly triggered by most TEs undergoing translation in Arabidopsis (Oberlin *et al*, 2022), it is possible that *EVD* hijacks the PTGS pathway to mediate copy number control (CNC) and regulate its own transposition rate. A variety of CNC factors have been found to be self-encoded by TEs, such as peptides to inhibit VLP formation or antisense ORFs to hamper reverse transcription, as means to minimize genomic damage on their host caused by over proliferation (Matsuda & Garfinkel, 2009; Cottee *et al*, 2021). However, *EVD* also reached high copy numbers in *nrpd1-* and *nrpe1-EVD* lines, despite the continuous action of PTGS. A full understanding of the impact of PTGS in *EVD* transposition and host recognition and defense mechanisms will require further investigation of *EVD* activity in genetic backgrounds defective for both PTGS and RdDM.

### PolIV-RdDM is essential for *EVD* control and *de novo* silencing

Recruitment of RdDM to *EVD-LTRs* independently of RDR6-RdDM, together with the absence of *EVD* TGS in either *nrpd1* or *nrpe1* backgrounds, indicates that PolIV-RdDM is essential for *de novo EVD* silencing during its genome colonization. Although absence of RdDM has little impact in the reactivation of silenced TEs in the Arabidopsis genome (He *et al*, 2021; 2022), a role for RdDM in controlling TE propagation has been previously shown for the heat-responsive LTR-retrotransposon (LTR-RTE) *ONSEN*, for which proliferation and copy number following induction is increased in *NRPD1* mutants (Ito *et al*, 2011; Matsunaga *et al*, 2012; Hayashi *et al*, 2020; Niu *et al*, 2022). Furthermore, genetic variation in the RdDM components *RDR2* and *NRPE1* has been associated with variation in TE content within Arabidopsis natural populations (Baduel *et al*, 2021; Sasaki *et al*, 2022; Jiang *et al*, 2023). Therefore, PolIV-RdDM might play a major role in *de novo* TE silencing besides its function as a DNA methylation maintenance pathway.

Coincidentally, RdDM at long, young, and potentially functional TEs, operates at their edges (Zemach *et al*, 2013; Stroud *et al*, 2013; 2014), similarly to the patterns found at *EVD* after the transition to TGS independently of PTGS. As mentioned previously, most LTR-RTEs intact enough to be translation-competent do not trigger PTGS when transcriptionally active (Oberlin *et al*, 2022). Hence, the phenomena of RdDM installation at flanking *LTRs* in absence of PTGS observed here for *EVD* might represent a more general mechanism of *de novo* silencing of LTR-RTEs.

### Potential mechanisms of TGS initiation in absence of PTGS

In absence of PTGS and the initial deposition of DNA methylation within the *EVD-GAG* sequence, the mechanism(s) triggering or initiating epigenetic silencing on *EVD* remain obscure. Apart from TE activity, several mechanisms have been shown to recruit RdDM or trigger the deposition of DNA methylation.

For example, double-stranded DNA breaks (DSBs) have been shown to trigger the production of siRNAs and promote the deposition of DNA methylation at the borders of the break points in Arabidopsis(Wei *et al*, 2012; Schalk *et al*, 2016; 2017; Du *et al*, 2022). In addition, little is known about the chromatin landscape of newly integrated TE copies, which might contain histone variants or histone modifications predisposing for DSB-induced silencing (Yelagandula *et al*, 2014; Lorković *et al*, 2017; Osakabe *et al*, 2018; Bourguet *et al*, 2021). However, such mechanism would imply that new insertions become individually silenced as they integrate, which is incompatible with the low levels of DNA methylation in lines with active *EVD* and its coordinated increase once the switch to TGS takes place. A significant number of simultaneous transposition events generating excessive DNA damage might be required to elicit a silencing response. Nonetheless, *EVD* does remain active in some individuals despite a high copy number and expression levels.

Given the arbitrary copy number at which TGS took place in *rdr6-EVD* lines, silencing might therefore depend on more stochastic events. Due to trans-activity of *EVD-LTR* 24-nt siRNAs, the initiation of RdDM in one or few *EVD* copies might suffice to spread silencing across all new active insertions. RNA hairpins resulting of TE tandem or nested insertions and acting as source of 24-nt siRNAs has been shown to mediate the epigenetic silencing of the *Mutator* TE in maize (Slotkin *et al*, 2003). Though this mechanism was previously deemed unlikely to trigger *EVD* TGS in *RDR6* wild type backgrounds (Marí-Ordóñez *et al*, 2013), increased transposition upon absence of PTGS might enhance the chances of hairpin formation. Nonetheless, RNA hairpins are generally processed into several siRNA sizes (Zilberman *et al*, 2004; Fusaro *et al*, 2006), and *LTR* siRNAs were predominantly 24-nt long.

Alternatively, and analogously to TE silencing triggered by transposition within piRNA clusters in *Drosophila melanogaster* (Brennecke *et al*, 2007; Goriaux *et al*, 2014; Guida *et al*, 2016), increased transposition might increase the probability of integration events within pre-existing heterochromatin domains, subjecting one or few *EVD* copies to RdDM. However, no increase in absolute or relative pericentromeric insertions was observed in *rdr6-EVD* lines that had switch to *EVD* TGS. A more detailed study with increased population size and better characterization of the chromatin environment in *EVD* landing sites will be required in the future to properly explore this possibility.

Additionally, the repetitiveness of *EVD* during a transposition burst, might led to changes in the three-dimensional chromatin organization forming, or bringing *EVD* to, chromatin interaction clusters associated with silencing (Grob *et al*, 2014; Grob & Grossniklaus, 2019). Furthermore, such repetitiveness might cause genome instability through non-allelic homologous recombination events (Sammarco *et al*, 2022) that trigger the silencing of *EVD*. Further investigation of the presence of genome rearrangements and chromatin conformation changes before and after *EVD* switch to TGS should help elucidate whether such changes are taking place and are indeed linked to *de novo EVD* silencing.

Finally, RdDM has been implicated in antiviral defense against Geminiviruses, plant single stranded circular DNA viruses replicating in the nucleus (Al-Kaff *et al*, 1998; Raja *et al*, 2008). Linear and circular extra chromosomal DNA (ecDNA and eccDNA respectively), products of reverse transcription, have been detected upon expression off several LTR-RTEs, including *EVD* (Lanciano *et al*, 2017; Mann *et al*, 2022; Niu *et al*, 2022). In absence of PTGS, *EVD* ecDNA and eccDNA content has been shown to increase in plant tissues (Lee *et al*, 2020; Zhang *et al*, 2023). Although it remains to be determined whether RdDM can act on TE ecDNA or eccDNA, they harbor the potential to become RdDM targets for the initiation of TGS on integrated *EVD* copies. In conclusion, our study reveals that PTGS alone is insufficient to explain the *de novo* initiation of epigenetic silencing of an active proliferative TE in the Arabidopsis genome and opens further questions about genome defense mechanisms. Several of the above-mentioned phenomena, alone or in combination, could contribute to *de novo* silencing of *EVD* and TEs, and deserve future investigation to gain insights into the mechanisms of defense against genomic parasites.

## Methods

### Plant material and growth conditions

Plants were grown on soil at 21 °C, in 16 hours light/ 8 hours dark cycle with an LED light intensity of 85 µM m^-2^ s^-1^. After germination, seedlings were transplanted at a rate of 1 plant per pot. Plants were genotyped by PCR as soon as a tip of a young rosette leaf could be harvested (See Supplementary Figure 8 for a list of primers used). Inflorescences (closed flower buds, no visible petals) were harvested, flash frozen in liquid nitrogen and stored at -70 °C. Cauline leaves used for *EVD* copy number quantification were harvested on ice and immediately stored at -70 °C. *rdr6-15* (NASC ID: N87978) was previously described in (Fahlgren *et al*, 2006); *nrpd1a-4* (*nrpd1*, NASC ID: N583051) in and *nrpd1b-11* (*nrpe1*, NASC ID: N529919) were described in (Alonso *et al*, 2003; Pontier *et al*, 2005). *rdr6-15* lines crossed to epi15 F8 were described in (Oberlin *et al*, 2022) and epi15 F11 were described in (Reinders *et al*, 2009; Marí-Ordóñez *et al*, 2013).

### Estimation of *EVD* expression and copy number by qPCR

Total RNA was extracted from 6-8 closed inflorescences ground to a fine powder by TRIzol™ (Thermo Fisher, #15596018) according to manufacturer’s instructions and precipitated in 1 volume of cold isopropanol. Total RNA was treated with RNase-free DNase I (Thermo Fisher #EN0521) and cDNA synthesis performed using the RevertAid Reverse Transcription Kit with random hexamer primers (Thermo Fisher #K1622). qPCR reactions were performed in a total volume of 10 μl using the KAPA SYBR FAST qPCR Master Mix (2X) (Sigma Aldrich #KK4602) with low ROX reference dye according to the manufacturer’s instruction and run on a QuantStudio 5 Real-Time PCR System (Thermo Fisher). Each reaction was performed in 2-3 technical. Relative expression was calculated as fold change of the ratio of target of interest and ACT2 (AT3G18780). For *EVD* copy number estimations, DNA was extracted from frozen ground cauline leaves using Edwards buffer (200 mM Tris pH 8, 200 mM NaCl, 25 mM EDTA, 0,5% (v/v) SDS). Samples were precipitated in 1 volume of 70% ethanol and diluted at 1/100 in double-distilled water. *EVD* copy number was estimated by absolute quantification from the ΔΔCt EVD and ACT2 levels, normalized by their inherent copy numbers of two in WT plants, respectively. Oligonucleotides used are listed in Supplementary Figure 8. Data analysis was done in Excel (Microsoft) and R (R Core Team (2023), R Foundation for Statistical Computing, Vienna, Austria, https://www.R-project.org). Plots were generated using ggplot2 (Wickham, 2016). Statistical significance was calculated using pairwise Student’s t-test.

### Small RNA blot analysis

Total RNA was extracted from 6-8 closed inflorescences ground to a fine powder by TRIzol™ (Thermo Fisher, #15596018) according to manufacturer’s instructions and precipitated in 1 volume of cold isopropanol. 10-20 μg of total RNA were mixed with an equal volume of 2X Novex^TM^ TBE-Urea sample buffer (Thermo Fisher #LLC6876) and resolved on a 17.5% polyacrylamide-ureal gel according to manufacturers instructions (National Diagnostics #EC-830, EC-835, EC-840), followed by electroblotting on a Hybond-NX Nylon membrane (Cytivia #RPN303T) and chemical crosslinking (12.5M 1-methylimidazol (Merk, #M50834), 31.25mg/mL N-(3-dimethylaminopropyl)-N’-ethylcarbodiimide hydrochloride (Merk, #E7750)) according to (Pall & Hamilton, 2008). PCR probes were labelled with [α-^32^P]-dCTP (Hartmann Analytic #SRP-305) using the Prime-it II Random Primer Labelling Kit (Agilent, #EK0031) and purified on illustra Microspin G-50 columns (Cytiva #27533001) according to manufacturers instructions. Oligo probes were labelled using [𝛾-^32^P]-ATP (Hartmann analytic #SRP-501), using T4 Polynucleotide kinase (Thermofisher, #EK0031) according to manufacturers instructions and purified on illustra Microspin G25 columns (Cytiva #27532501). Hybridization was performed in PerfectHyb hybridization buffer (Sigma Aldricht #H7033) for PCR probes, or in Church buffer (7% SDS, 0.5 M Na2HPO4/NaH2PO4 pH 7.2, 1mM EDTA) for oligonucleotide probes. Following over-night hybridization at 42°C in a rotary oven, membranes were washed three times at 50°C with 2XSCC, 0.1% (v/v) SDS. Detection was performed with a Typhoon FLA 9500 (GE Healthcare). Oligonucleotide used for probes are listen in Supplementary Figure 8.

### Methylation analysis by EM-Seq

DNA was extracted from closed inflorescences from single plants, using the DNeasy plant kit (Qiagen, #69204) according to manufacturer’s instructions. Libraries were prepared using the NEBNext® Enzymatic Methyl-seq Kit (NEB, #M7634) following the manufacturer’s instructions. Sequencing was performed on an Illumina NovaSeq SP to produce paired-end reads of 150 bp. Sequenced reads were quality filtered and adaptor trimmed using TrimGalore version 0.6.2 (https://github.com/FelixKrueger/TrimGalore). Enzymatic-converted reads were aligned to the TAIR10 Arabiopsis genome using Bismark version v0.24.2 with bismark (--unmapped --ambiguous --maxins 700) (Krueger & Andrews, 2011) PCR/optical duplicates were removed using *deduplicate_bismark* and the resulting BAM files were further processed to generate cytosine files *bismark_methylation_extracto*r (--paired-end --cytosine_report --CX_context --comprehensive --no_overlap). Weighted average methylation (Schultz *et al*, 2012) was calculated and for cytonises laying on the each *EVD* domain. Conversion rates were assessed by calculating the average methylation level for reads mapping to the unmethylated chloroplast (Cokus *et al*, 2008). Statistical testing was done in R using the function wilcox_test from the rstatix package. P-values were adjusted for multiple testing using Benjamini & Hochberg. Meaning of significance : "****" : < 10^-4^, "***" : < 10-^3^, "**" : < 0.01, "*" : < 0.05, "ns": ≥ 0.05.

#### *EVD* copy number estimation and mapping of new insertions from EM-seq. Coverage-based copy number estimation

Depth of coverage at every base between the 5’UTR and 3’UTR of reference EVD insertion (using your annotation Arturo) (Chr5:5630399-5634902), as well as on two ∼10kb intervals upstream and downstream of EVD (Chr5:5619977-5629277, Chr5:5636011-5645311) was calculated for each paired-end BAM using *samtools depth* (-aa) (Li *et al*, 2009) (https://github.com/samtools/samtools). The ratio between mean depth of coverage on EVD and mean depth of coverage on the upstream and downstream intervals was calculated for each sample. The mean inverse ratio from Col-0 and *rdr6-15* samples only carrying the reference insertion were used as a normalization factor in every other sample.

#### Discordant read mate pairs-based mapping of *EVD* insertions

For the identification of new *EVD* insertions, an existing protocol for the detection of TE insertions from short-read sequenceing data, was adapted for EM-seq datasets (Baduel *et al*, 2021). Each pair of fastq files containing unmapped mates was remapped in single-end (-s) mode using *bismark* (--local for both mates, --pbat for second in pair) to the TAIR10 genome, as well as to the EVD DNA sequence extracted from TAIR10 (Chr5:5629976-5635312) with parameters --local -D 20 -R 3 -N 1 for both mates, --pbat for second in pair. Since multi-mapping reads are automatically removed by *bismark*, part of the 3’ LTR from the extracted EVD sequence was masked using *bedtools maskfasta* (Chr5:5635050:5635156) to prevent reads from mapping exactly to both LTRs, leading to some mismapped reads but maximizing coverage. Fragments for which one mate mapped onto the extracted EVD sequence and the other to any other position in the genome were considered as discordant and candidate reads for identification of de novo insertions. Cases where a single read would map to both EVD and another genomic location were discarded.

Candidate reads were extracted using *bedtools bamtobed*, and loaded into R (v4.2.0) (https://www.r-project.org) for processing. A simple scheme was used to identify potential insertions: reads that mapped outside of EVD were sorted by chromosome and start coordinates, and then put into clusters based on their proximity. The directionality (relative to the reference genome) of their component reads was deduced from their strandness and their pairing (first or second in pair). The directionality of candidate reads mapping around a bona fide non-reference insertion is known as they should all point towards the insertion site. Therefore, valid clusters (comprised of *m+n* reads) where considered those for which the first *m* reads (upstream of the insertion site) pointed towards the 3’ end of the reference chromosome and the remaining *n* reads (downstream of the insertion site) pointed towards the 5’ end of the reference chromosome (m >= 3, n >= 3), from those the insertion site was inferred as located between the end coordinate of the *mth* read and the start coordinates of the *m+1st* read. Similarly, paired-end mates of the first *m* reads (or last *n*) should all share the same directionality and additionally, the first *m* and the last *n* reads display opposite directionality. Any cluster that did not adhere to these rules was discarded. Within each cluster, the directionality of reads mapping onto *EVD* was used to asses the strandness of the insertion, and from it each read’s LTR-of-origin inferred.

For each remaining cluster, presence and size of target-site-duplicatiion (TSD) was established by inspecting the cigar strings of the *mth* and *m+1st* reads. If the *mth* read was soft-clipped on its 5’ end and the *m+1st* read was soft-clipped on its 3’ end, and both reads overlapped each other (which was always true), then a TSD was deemed present and the size of the overlap taken as the size of the TSD. 91% of clusters showed such a pattern.

### Assessment of DNA methylation levels in new *EVD* insertions

For each identified new insertion, the associated cluster of reads was retrieved and LTR-of-origin determined individually for every read. In each sample, the appropriate BAM file was subsampled so it would only include the appropriate clustered reads containing LTR-of-origing information using *picard FilterSamReads* (https://broadinstitute.github.io/picard/). Por each sample, cluster and LTR-of-origin, a cytosine report was obtained from the subsampled BAM as specified before and a single methylation level for each cytosine context was obtained using weighted average methylation as described above was calculated and for cytonises laying on the each individual LTR.

### Bisulfite-PCR sequencing based DNA methylation analysis

DNA was extracted from 6-8 closed inflorescences bulked from 4 plants ground to a fine powder, using the DNeasy plant kit (Qiagen, #69204) according to manufacturer’s instructions. Bisulfite treatment was performed with the EZ DNA Methylation-Gold Kit (Zymo Research, #D5005), using the manufacturer’s standard protocol. DNA fragments of interest were amplified by PCR. PCR reactions and primer design was conducted according to published recommendations (Supplementary Figure 8, (Henderson *et al*, 2010)) and cloned using CloneJET PCR Cloning Kit (Thermo, #K1231). Single colonies were sequenced by Sanger sequencing. Sequences were analysed using the browser based Kismeth software (https://katahdin.girihlet.com/kismeth/revpage.pl) (Gruntman *et al*, 2008). Wilson score intervals were used to find 95% confidence intervals for the percentages of methylated sites in each context. Results shown were obtained from two independent experiments.

## Data availability

An inventory of all the raw data used to generate each figure panel (as well as those in Supplementary information) along with all raw images, qPCR and sanger sequencing files, can be found in Zenodo (www.zenodo.org) under the doi: 10.5281/zenodo.11350776. NGS data generated for this study are accessible at NCBI SRA under ID number PRJNA1111825, submission ID SUB14423611.

## Author contributions

M.T. and A.M-O conceived and designed the study. M.T carried most experiments together with V.B-B and assisted by L.D-N. G.B-H performed EM-seq computational analysis. L.L. conducted DNA methylation analysis by BS-PCR. M.T. and G.B-H performed statistical analysis. A.M-O and M.T interpreted the results and wrote the manuscript. All authors have read and validated the manuscript.

## Acknowledgements

This work was supported by the Gregor Mendel Institute of the Austrian Academy of Sciences core funding attributed to A.M-O and M.N. We thank current and former members of the Marí-Ordóñez group as well as colleagues from the Gregor Mendel Institute and the Vienna BioCenter campus for useful discussions, ideas and feedback on the project and manuscript. We are grateful to the Vienna BioCenter Core Facilities (VBCF): NGS facility for EM-seq library preparation and sequencing, and the Plant Science Facility for assistance with plant work. We also thank the IMP Molecular Biology Service for providing reagents and Sanger sequencing. Lastly, we would like to acknowledge the VBC in-house COVID-19 testing facility for their efforts in providing a safe working environment during the pandemic.

## References

Al-Kaff NS, Covey SN, Kreike MM, Page AM, Pinder R & Dale PJ (1998) Transcriptional and posttranscriptional plant gene silencing in response to a pathogen. Science 279: 2113–2115

Allshire RC & Madhani HD (2018) Ten principles of heterochromatin formation and function. Nat Rev Mol Cell Biol 19: 229–244

Alonso JM, Stepanova AN, Leisse TJ, Kim CJ, Chen H, Shinn P, Stevenson DK, Zimmerman J, Barajas P, Cheuk R, et al (2003) Genome-Wide Insertional Mutagenesis of Arabidopsis thaliana. Science 301: 653–657

Baduel P, Leduque B, Ignace A, Gy I, Gil J, Loudet O, Colot V & Quadrana L (2021) Genetic and environmental modulation of transposition shapes the evolutionary potential of Arabidopsis thaliana. Genome Biol 22: 138

Baduel P, Quadrana L & Colot V (2021) Plant Transposable Elements, Methods and Protocols. Methods Mol Biol 2250: 157–169

Bennetzen JL & Wang H (2014) The Contributions of Transposable Elements to the Structure, Function, and Evolution of Plant Genomes. Plant Biology 65: 505–530

Bernatavichute YV, Zhang X, Cokus S, Pellegrini M & Jacobsen SE (2008) Genome-Wide Association of Histone H3 Lysine Nine Methylation with CHG DNA Methylation in Arabidopsis thaliana. PLoS ONE 3: e3156

Bourguet P, Picard CL, Yelagandula R, Pélissier T, Lorković ZJ, Feng S, Pouch-Pélissier M-N, Schmücker A, Jacobsen SE, Berger F, et al (2021) The histone variant H2A.W and linker histone H1 co-regulate heterochromatin accessibility and DNA methylation. Nat Commun 12: 2683

Bourque G, Burns KH, Gehring M, Gorbunova V, Seluanov A, Hammell M, Imbeault M, Izsvák Z, Levin HL, Macfarlan TS, et al (2018) Ten things you should know about transposable elements. Genome biology 19: 1– 12

Brennecke J, Aravin AA, Stark A, Dus M, Kellis M, Sachidanandam R & Hannon GJ (2007) Discrete small RNA-generating loci as master regulators of transposon activity in Drosophila. Cell 128: 1089–1103

Chan SWL, Henderson IR & Jacobsen SE (2005) Gardening the genome: DNA methylation in Arabidopsis thaliana. Nature Reviews Genetics 6: 351–360

Cokus SJ, Feng S, Zhang X, Chen Z, Merriman B, Haudenschild CD, Pradhan S, Nelson SF, Pellegrini M & Jacobsen SE (2008) Shotgun bisulphite sequencing of the *Arabidopsis* genome reveals DNA methylation patterning. Nature 452: 215–219

Cottee MA, Beckwith SL, Letham SC, Kim SJ, Young GR, Stoye JP, Garfinkel DJ & Taylor IA (2021) Structure of a Ty1 restriction factor reveals the molecular basis of transposition copy number control. Nat Commun 12: 5590

Du J, Johnson LM, Groth M, Feng S, Hale CJ, Li S, Vashisht AA, Gallego-Bartolome J, Wohlschlegel JA, Patel DJ, et al (2014) Mechanism of DNA Methylation-Directed Histone Methylation by KRYPTONITE. Mol Cell 55: 495– 504

Du J, Liu Y, Lu L, Shi J, Xu L, Li Q, Cheng X, Chen J & Zhang X (2022) Accumulation of DNA damage alters microRNA gene transcription in Arabidopsis thaliana. Bmc Plant Biol 22: 576

Du J, Zhong X, Bernatavichute YV, Stroud H, Feng S, Caro E, Vashisht AA, Terragni J, Chin HG, Tu A, et al (2012) Dual Binding of Chromomethylase Domains to H3K9me2-Containing Nucleosomes Directs DNA Methylation in Plants. Cell 151: 167–180

Erdmann RM & Picard CL (2020) RNA-directed DNA Methylation. Plos Genet 16: e1009034

Fahlgren N, Montgomery TA, Howell MD, Allen E, Dvorak SK, Alexander AL & Carrington JC (2006) Regulation of AUXIN RESPONSE FACTOR3 by TAS3 ta-siRNA Affects Developmental Timing and Patterning in Arabidopsis. Curr Biol 16: 939–944

Feng S, Zhong Z, Wang M & Jacobsen SE (2020) Efficient and accurate determination of genome-wide DNA methylation patterns in Arabidopsis thaliana with enzymatic methyl sequencing. Epigenetics & chromatin 13: 42–17

Fusaro AF, Matthew L, Smith NA, Curtin SJ, Dedic-Hagan J, Ellacott GA, Watson JM, Wang M, Brosnan C, Carroll BJ, et al (2006) RNA interference-inducing hairpin RNAs in plants act through the viral defence pathway. EMBO Rep 7: 1168–1175

Gilly A, Etcheverry M, Madoui M-A, Guy J, Quadrana L, Alberti A, Martin A, Heitkam T, Engelen S, Labadie K, et al (2014) TE-Tracker: systematic identification of transposition events through whole-genome resequencing. BMC Bioinformatics 15: 377

Goriaux C, Théron E, Brasset E & Vaury C (2014) History of the discovery of a master locus producing piRNAs: the flamenco/COM locus in Drosophila melanogaster. Frontiers Genetics 5: 257

Grob S & Grossniklaus U (2019) Invasive DNA elements modify the nuclear architecture of their insertion site by KNOT-linked silencing in Arabidopsis thaliana. 1–15

Grob S, Schmid MW & Grossniklaus U (2014) Hi-C Analysis in Arabidopsis Identifies the KNOT, a Structure with Similarities to the flamenco Locus of Drosophila. Molecular Cell 55: 678–693

Gruntman E, Qi Y, Slotkin RK, Roeder T, Martienssen RA & Sachidanandam R (2008) Kismeth: Analyzer of plant methylation states through bisulfite sequencing. Bmc Bioinformatics 9: 371

Guida V, Cernilogar FM, Filograna A, Gregorio RD, Ishizu H, Siomi MC, Schotta G, Bellenchi GC & Andrenacci D (2016) Production of Small Noncoding RNAs from the flamenco Locus Is Regulated by the gypsy Retrotransposon of Drosophila melanogaster. Genetics 204: 631–644

Hayashi Y, Takehira K, Nozawa K, Suzuki T, Masuta Y, Kato A & Ito H (2020) ONSEN shows different transposition activities in RdDM pathway mutants. Genes & genetic systems: 20–00019

He L, Huang H, Bradai M, Zhao C, You Y, Ma J, Zhao L, Lozano-Durán R & Zhu J-K (2022) DNA methylation-free Arabidopsis reveals crucial roles of DNA methylation in regulating gene expression and development. Nat Commun 13: 1335

He L, Zhao C, Zhang Q, Zinta G, Wang D, Lozano-Durán R & Zhu J-K (2021) Pathway conversion enables a double-lock mechanism to maintain DNA methylation and genome stability. Proc Natl Acad Sci 118: e2107320118

Henderson IR, Chan SR, Cao X, Johnson L & Jacobsen SE (2010) Accurate sodium bisulfite sequencing in plants. Epigenetics 5: 47–49

Ito H, Gaubert H, Bucher E, Mirouze M, Vaillant I & Paszkowski J (2011) An siRNA pathway prevents transgenerational retrotransposition in plants subjected to stress. Nature 472: 115–119

Jedlicka P, Lexa M, Vanat I, Hobza R & Kejnovsky E (2019) Nested plant LTR retrotransposons target specific regions of other elements, while all LTR retrotransposons often target palindromes and nucleosome-occupied regions: in silico study. Mob DNA 10: 50

Jiang J, Xu Y-C, Zhang Z-Q, Chen J-F, Niu X-M, Hou X-H, Li X-T, Wang L, Zhang YE, Ge S, et al (2023) Forces driving transposable element load variation during Arabidopsis range expansion. Plant Cell 36: 840–862

Krueger F & Andrews SR (2011) Bismark: a flexible aligner and methylation caller for Bisulfite-Seq applications. Bioinformatics 27: 1571–1572

Kuhlmann M & Mette MF (2012) Developmentally non-redundant SET domain proteins SUVH2 and SUVH9 are required for transcriptional gene silencing in Arabidopsis thaliana. Plant Mol Biol 79: 623–633

Lanciano S, Carpentier M-C, Llauro C, Jobet E, Robakowska-Hyzorek D, Lasserre E, Ghesquière A, Panaud O & Mirouze M (2017) Sequencing the extrachromosomal circular mobilome reveals retrotransposon activity in plants. Plos Genet 13: e1006630

Law JA, Du J, Hale CJ, Feng S, Krajewski K, Palanca AMS, Strahl BD, Patel DJ & Jacobsen SE (2013) Polymerase IV occupancy at RNA-directed DNA methylation sites requires SHH1. Nature Publishing Group 498: 385–389

Law JA & Jacobsen SE (2010) Establishing, maintaining and modifying DNA methylation patterns in plants and animals. Nature Reviews Genetics 11: 204–220

Lee SC, Ernst E, Berube B, Borges F, Parent J-S, Ledon P, Schorn A & Martienssen RA (2020) Arabidopsis retrotransposon virus-like particles and their regulation by epigenetically activated small RNA. Genome Research 30: 576–588

Lee Y-S, Maple R, Dürr J, Dawson A, Tamim S, Genio C del, Papareddy R, Luo A, Lamb JC, Amantia S, et al (2021) A transposon surveillance mechanism that safeguards plant male fertility during stress. Nat Plants 7: 34–41

Li H, Handsaker B, Wysoker A, Fennell T, Ruan J, Homer N, Marth G, Abecasis G, Durbin R & Subgroup 1000 Genome Project Data Processing (2009) The Sequence Alignment/Map format and SAMtools. Bioinformatics 25: 2078–2079

Lippman Z, Gendrel A-V, Black M, Vaughn MW, Dedhia N, McCombie WR, Lavine K, Mittal V, May B, Kasschau KD, et al (2004) Role of transposable elements in heterochromatin and epigenetic control. Nature 430: 471– 476

Lorković ZJ, Park C, Goiser M, Jiang D, Kurzbauer M-T, Schlögelhofer P & Berger F (2017) Compartmentalization of DNA Damage Response between Heterochromatin and Euchromatin Is Mediated by Distinct H2A Histone Variants. Current Biology 27: 1192–1199

Mann L, Seibt KM, Weber B & Heitkam T (2022) ECCsplorer: a pipeline to detect extrachromosomal circular DNA (eccDNA) from next-generation sequencing data. Bmc Bioinformatics 23: 40

Marí-Ordóñez A, Marchais A, Etcheverry M, Martin A, Colot V & Voinnet O (2013) Reconstructing de novo silencing of an active plant retrotransposon. Nature Genetics 45: 1029–1039

Mathieu O, Reinders J, Čaikovski M, Smathajitt C & Paszkowski J (2007) Transgenerational Stability of the Arabidopsis Epigenome Is Coordinated by CG Methylation. Cell 130: 851–862

Matsuda E & Garfinkel DJ (2009) Posttranslational interference of Ty1 retrotransposition by antisense RNAs. Proc Natl Acad Sci 106: 15657–15662

Matsunaga W, Kobayashi A, Kato A & Ito H (2012) The effects of heat induction and the siRNA biogenesis pathway on the transgenerational transposition of ONSEN, a copia-like retrotransposon in Arabidopsis thaliana. Plant and Cell Physiology 53: 824–833

Matzke MA & Mosher RA (2014) RNA-directed DNA methylation: an epigenetic pathway of increasing complexity. Nature Reviews Genetics 15: 394–408

McClintock B (1984) The Significance of Responses of the Genome to Challenge. Science 226: 792–801

McCue AD, Panda K, Nuthikattu S, Choudury SG, Thomas EN & Slotkin RK (2015) ARGONAUTE 6 bridges transposable element mRNA-derived siRNAs to the establishment of DNA methylation. The EMBO journal 34: 20–35

Mirouze M, Reinders J, Bucher E, Nishimura T, Schneeberger K, Ossowski S, Cao J, Weigel D, Paszkowski J & Mathieu O (2009) Selective epigenetic control of retrotransposition in Arabidopsis. Nature 461: 427–430

Niu X, Chen L, Kato A & Ito H (2022) Regulatory mechanism of a heat-activated retrotransposon by DDR complex in Arabidopsis thaliana. Front Plant Sci 13: 1048957

Nuthikattu S, McCue AD, Panda K, Fultz D, DeFraia C, Thomas EN & Slotkin RK (2013) The Initiation of Epigenetic Silencing of Active Transposable Elements Is Triggered by RDR6 and 21-22 Nucleotide Small Interfering RNAs. PLANT PHYSIOLOGY 162: 116–131

Oberlin S, Rajeswaran R, Trasser M, Barragán-Borrero V, Schon MA, Plotnikova A, Loncsek L, Nodine MD, Marí-Ordóñez A & Voinnet O (2022) Innate, translation-dependent silencing of an invasive transposon in Arabidopsis. Embo Rep 23: e53400

Oberlin S, Sarazin A, Chevalier C, Voinnet O & Marí-Ordóñez A (2017) A genome-wide transcriptome and translatome analysis of Arabidopsistransposons identifies a unique and conserved genome expression strategy for Ty1/Copiaretroelements. Genome Research 27: 1549–1562

Orozco-Arias S, Isaza G & Guyot R (2019) Retrotransposons in Plant Genomes: Structure, Identification, and Classification through Bioinformatics and Machine Learning. Int J Mol Sci 20: 3837

Osakabe A, Lorković ZJ, Kobayashi W, Tachiwana H, Yelagandula R, Kurumizaka H & Berger F (2018) Histone H2A variants confer specific properties to nucleosomes and impact on chromatin accessibility. Nucleic Acids Research 46: 7675–7685

Pall GS & Hamilton AJ (2008) Improved northern blot method for enhanced detection of small RNA. Nat Protoc 3: 1077–1084

Panda K, Ji L, Neumann DA, Daron J, Schmitz RJ & Slotkin RK (2016) Full-length autonomous transposable elements are preferentially targeted by expression-dependent forms of RNA-directed DNA methylation. Genome Biology 17: 170–19

Pontier D, Yahubyan G, Vega D, Bulski A, Saez-Vasquez J, Hakimi M-A, Lerbs-Mache S, Colot V & Lagrange T (2005) Reinforcement of silencing at transposons and highly repeated sequences requires the concerted action of two distinct RNA polymerases IV in Arabidopsis. Gene Dev 19: 2030–2040

Quadrana L, Etcheverry M, Gilly A, Caillieux E, Madoui M-A, Guy J, Silveira AB, Engelen S, Baillet V, Wincker P, et al (2019) Transposition favors the generation of large effect mutations that may facilitate rapid adaption. Nat Commun 10: 3421–10

Quadrana L, Silveira AB, Mayhew GF, LeBlanc C, Martienssen RA, Jeddeloh JA & Colot V (2016) The Arabidopsis thaliana mobilome and its impact at the species level. Elife 5

Raja P, Sanville BC, Buchmann RC & Bisaro DM (2008) Viral Genome Methylation as an Epigenetic Defense against Geminiviruses. J Virol 82: 8997–9007

Reinders J, Wulff BBH, Mirouze M, Ordóñez AM, Dapp M, Rozhon W, Bucher E, Theiler G & Paszkowski J (2009) Compromised stability of DNA methylation and transposon immobilization in mosaic Arabidopsis epigenomes. Genes & development 23: 939–950

Roquis D, Robertson M, Yu L, Thieme M, Julkowska M & Bucher E (2021) Genomic impact of stress-induced transposable element mobility in Arabidopsis. Nucleic Acids Res 49: gkab828-

Sammarco I, Pieters J, Salony S, Toman I, Zolotarov G & Placette CL (2022) Epigenetic targeting of transposon relics: beating the dead horses of the genome? Epigenetics 17: 1331–1344

Sasaki E, Gunis J, Reichardt-Gomez I, Nizhynska V & Nordborg M (2022) Conditional GWAS of non-CG transposon methylation in Arabidopsis thaliana reveals major polymorphisms in five genes. PLoS Genet 18: e1010345

Schalk C, Cognat V, Graindorge S, Vincent T, Voinnet O & Molinier J (2017) Small RNA-mediated repair of UV-induced DNA lesions by the DNA DAMAGE-BINDING PROTEIN 2 and ARGONAUTE 1. Proc National Acad Sci 114: E2965–E2974

Schalk C, Drevensek S, Kramdi A, Kassam M, Ahmed I, Cognat V, Graindorge S, Bergdoll M, Baumberger N, Heintz D, et al (2016) DNA DAMAGE BINDING PROTEIN2 Shapes the DNA Methylation Landscape. Plant Cell 28: 2043–2059

Schubert I & Vu GTH (2016) Genome Stability and Evolution: Attempting a Holistic View. Trends Plant Sci 21: 749– 757

Schultz MD, Schmitz RJ & Ecker JR (2012) ‘Leveling’ the playing field for analyses of single-base resolution DNA methylomes. Trends Genet 28: 583–585

Sigman MJ, Panda K, Kirchner R, McLain LL, Payne H, Peasari JR, Husbands AY, Slotkin RK & McCue AD (2021) An siRNA-guided ARGONAUTE protein directs RNA polymerase V to initiate DNA methylation. Nat Plants 7: 1461–1474

Slotkin RK, Freeling M & Lisch D (2003) Mu killer Causes the Heritable Inactivation of the Mutator Family of Transposable Elements in Zea mays. Genetics 165: 781–797

Slotkin RK, Vaughn M, Borges F, TanurdZiC M, Becker JoD, FeijO JA & Martienssen RA (2009) Epigenetic Reprogramming and Small RNA Silencing of Transposable Elements in Pollen. Cell 136: 461–472

Stroud H, Do T, Du J, Zhong X, Feng S, Johnson L, Patel DJ & Jacobsen SE (2014) Non-CG methylation patterns shape the epigenetic landscape in Arabidopsis. Nat Struct Mol Biol 21: 64–72

Stroud H, Greenberg MVC, Feng S, Bernatavichute YV & Jacobsen SE (2013) Comprehensive Analysis of Silencing Mutants Reveals Complex Regulation of the Arabidopsis Methylome. Cell 152: 352–364

Stuart T, Eichten SR, Cahn J, Karpievitch YV, Borevitz JO & Lister R (2016) Population scale mapping of transposable element diversity reveals links to gene regulation and epigenomic variation. eLife 5: e20777

Taochy C, Yu A, Bouché N, Bouteiller N, Elmayan T, Dressel U, Carroll BJ & Vaucheret H (2019) Post-transcriptional gene silencing triggers dispensable DNA methylation in gene body in Arabidopsis. Nucleic Acids Res 47: 9104–9114

Teixeira FK, Heredia F, Sarazin A, Roudier F, Boccara M, Ciaudo C, Cruaud C, Poulain J, Berdasco M, Fraga MF, et al (2009) A Role for RNAi in the Selective Correction of DNA Methylation Defects. Science 323: 1600–1604

Thieme M, Lanciano S, Balzergue S, Daccord N, Mirouze M & Bucher E (2017) Inhibition of RNA polymerase II allows controlled mobilisation of retrotransposons for plant breeding. Genome Biol 18: 134

Tsukahara S, Kobayashi A, Kawabe A, Mathieu O, Miura A & Kakutani T (2009) Bursts of retrotransposition reproduced in Arabidopsis. Nature: 1–5

Wei W, Ba Z, Gao M, Wu Y, Ma Y, Amiard S, White CI, Danielsen JMR, Yang Y-G & Qi Y (2012) A Role for Small RNAs in DNA Double-Strand Break Repair. Cell 149: 101–112

Wickham H (2016) ggplot2, Elegant Graphics for Data Analysis. R: 189–201

Wu L, Mao L & Qi Y (2012) Roles of dicer-like and argonaute proteins in TAS-derived small interfering RNA-triggered DNA methylation. Plant Physiol 160: 990–999

Yelagandula R, Stroud H, Holec S, Zhou K, Feng S, Zhong X, Muthurajan UM, Nie X, Kawashima T, Groth M, et al (2014) The Histone Variant H2A.W Defines Heterochromatin and Promotes Chromatin Condensation in Arabidopsis. Cell 158: 98–109

Yoon S, Xuan Z, Makarov V, Ye K & Sebat J (2009) Sensitive and accurate detection of copy number variants using read depth of coverage. Genome Res 19: 1586–1592

Zemach A, Kim MY, Hsieh P-H, Coleman-Derr D, Eshed-Williams L, Thao K, Harmer SL & Zilberman D (2013) The Arabidopsis Nucleosome Remodeler DDM1 Allows DNA Methyltransferases to Access H1-Containing Heterochromatin. Cell 153: 193–205

Zhang P, Mbodj A, Soundiramourtty A, Llauro C, Ghesquière A, Ingouff M, Slotkin RK, Pontvianne F, Catoni M & Mirouze M (2023) Extrachromosomal circular DNA and structural variants highlight genome instability in Arabidopsis epigenetic mutants. Nat Commun 14: 5236

Zilberman D, Cao X, Johansen LK, Xie Z, Carrington JC & Jacobsen SE (2004) Role of Arabidopsis ARGONAUTE4 in RNA-Directed DNA Methylation Triggered by Inverted Repeats. Curr Biol 14: 1214–1220

