## Supplementary information for "PTGS is dispensable for the initiation of epigenetic silencing of an active transposon in *Arabidopsis*"

#### Table of content:

Page

2 **Supplementary Figure 1:** *EVD GAG-POL* expression in *RDR6* wild-type and mutant backgrounds.
3 **Supplementary Figure 2:** EM-seq general stats.
4 **Supplementary Figure 3:** *EVD-POL* and *3'LTR* methylation levels inn *RDR6* wild-type and mutant backgrounds.
5 **Supplementary Figure 4:** *EVD* concordant and discordant paired read mates in EM-seq.
6 **Supplementary Figure 5:** Genomic positions of new *EVD* insertion in *RDR6- and rdr6-EVD* F6 individuals.
7 **Supplementary Figure 6:** BS-PCR analysis of *EVD-GAG* DNA methylation levels in RdDM mutants.
8 **Supplementary Figure 7**: BS-PCR analysis of *EVD-LTR* DNA methylation levels in RdDM mutants.
9 **Supplementary Figure 8:** Oligos used in this study.

### SUPPLEMENTARY FIGURE 1

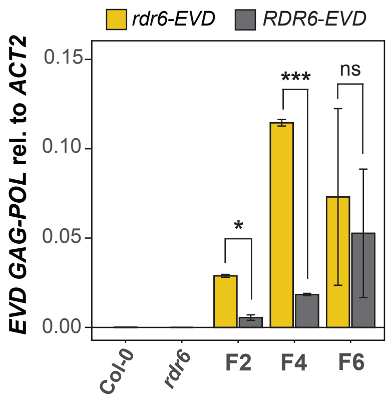


**Supplementary Figure 1: *EVD GAG-POL* expression in *RDR6* wild-type and mutant backgrounds.** qPCR analysis of *EVD flGAG-POL* expression normalized to *ACT2* in *EVD-RDR6* and *EVD-rdr6* lines at generations 2, 4 and 6. Two biological replicates are represented, error bars show standard error (n.s: not significant, $p>0.05$, *$p<0.05$, **$p<0.01$, ***$p<0.005$, two-sided t-test between indicated samples).

### SUPPLEMENTARY FIGURE 2

**A**

| sample | conversion rate |
|---|---|
| **Col-0 #1** | 99,857 % |
| **Col-0 #2** | 99,866 % |
| ***rdr6-15* #1** | 99,847 % |
| ***rdr6-15* #2** | 99,861 % |
| ***RDR6-EVD* #2** | 99,850 % |
| ***RDR6-EVD* #9** | 99,870 % |
| ***RDR6-EVD* #1** | 99,878 % |
| ***RDR6-EVD* #7** | 99,862 % |
| ***rdr6-EVD* #8** | 99,864 % |
| ***rdr6-EVD* #10** | 99,830 % |
| ***rdr6-EVD* #5** | 99,880 % |
| ***rdr6-EVD* #6** | 99,865 % |

**B**

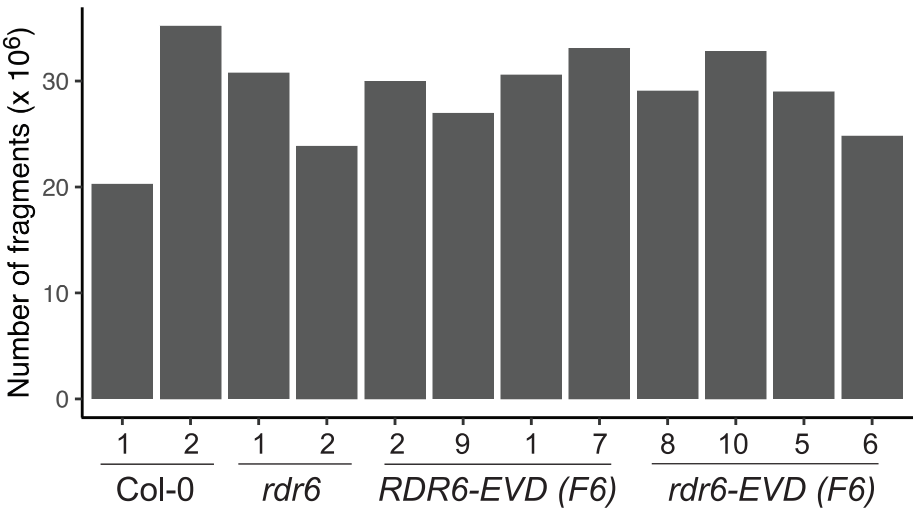


**C**

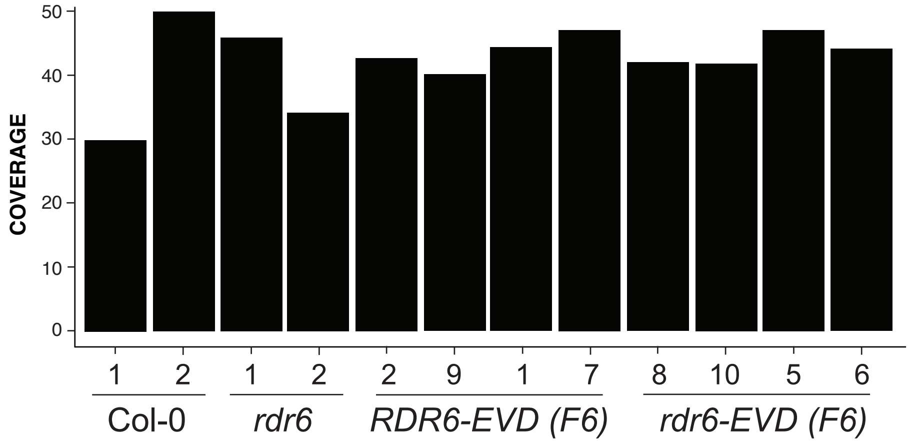


**Supplemenntary Figure 2: EM-seq general stats. (A)** Cytosine-to-thymine conversion rates of the unmethylated chloroplastic DNA for each EM-seq library (see materials and methods for further information). **(B)** Number of paired-end fragments obtained in each library. **(C)** Arabidopsis genome coverage in each library.

### SUPPLEMENTARY FIGURE 3

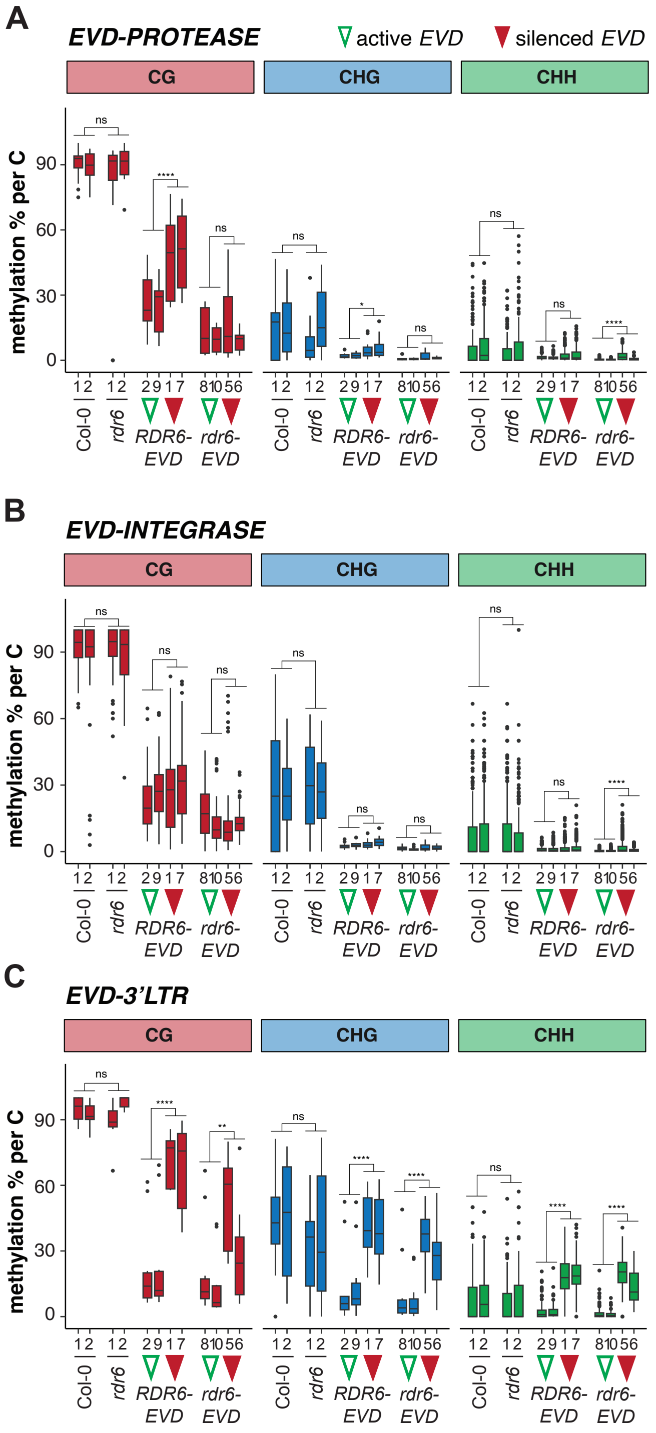


**Supplementary Figure 3: *EVD-POL* and *3'LTR* methylation levels in *RDR6* wild-type and mutant backgrounds.** DNA methylation % per cytosine for all three DNA methylation contexts in *EVD POL PROTEASE* **(A)**, *INTEGRASE* **(B)** and *3'LTR* domains **(C)** in WT, *rdr6, RDR6-EVD* and *rdr6-EVD* F6 lines with active and silenced *EVD* (empty green and filled red arrowheads respectively). In all boxplots: median is indicated by a solid bar, the boxes extend from the first to the third quartile and whiskers reach the furthest value within 1.5 times the interquartile range, dots indicate outliers outside of the above range. Wilcoxon rank sum test adjusted p-value between indicated groups of samples: ns: non significant ($p \geq 0.05$), *p<0.05, **p<0.01, ***p<0.001, ****p<0.0001.

### SUPPLEMENTARY FIGURE 4

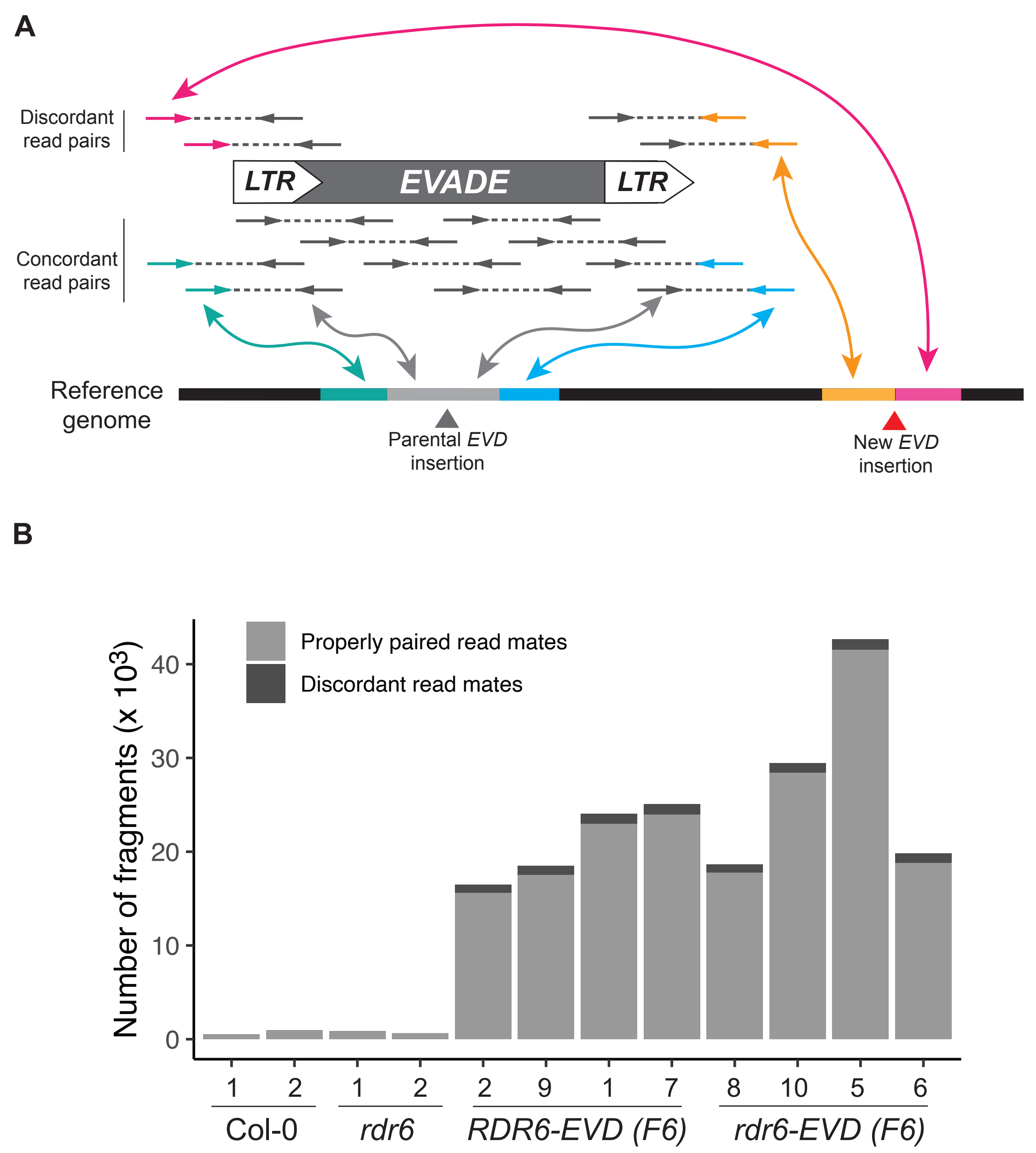


**Supplementary Figure 4: *EVD* concordant and discordant paired read mates in EM-seq. A** Schematic representation of the strategy used to map new EVD insertions using discordant read mates from EM-seq. **B** Number of fragments from concordant (properly paired) and discordant read mates mapping to *EVD* in each of the EM-seq libraries.

#### SUPPLEMENTARY FIGURE 5

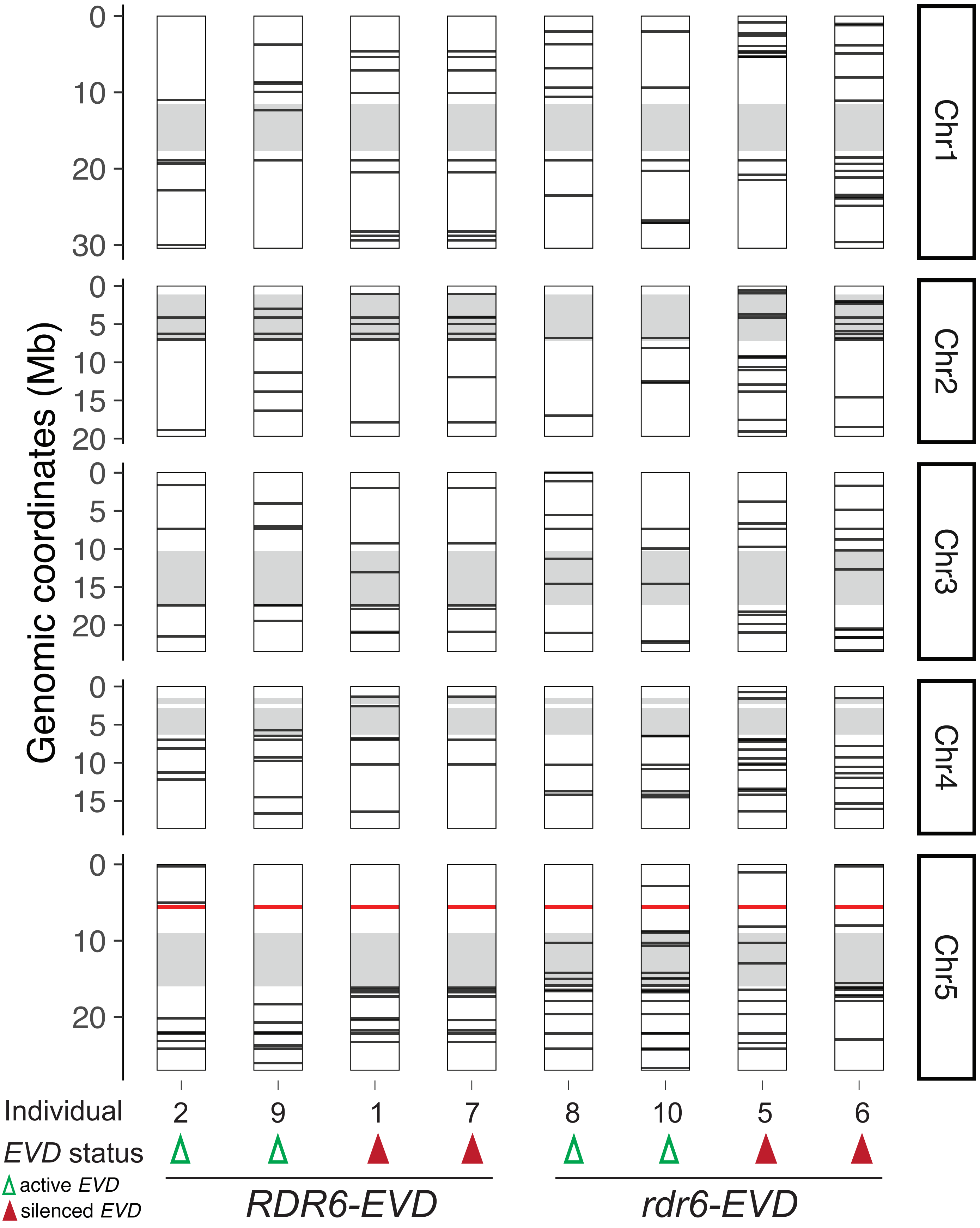


**Supplementary Figure 5: Genomic positions of new *EVD* insertions in *RDR6-* and *rdr6-EVD* F6 individuals.** Genomic location of new *EVD* insertions mapped through discordant read pair mates in EM-seq data (Figure 4, Supplementary Fig. 4) in the Arabidopsis genome. Parental *EVD* (AT5G17125) location is indicated with a red line. New *EVD* insertions are marked with black lines. Pericentromeric regions in each of Arabidopsis five chromosomes are marked in grey.

### SUPPLEMENTARY FIGURE 6

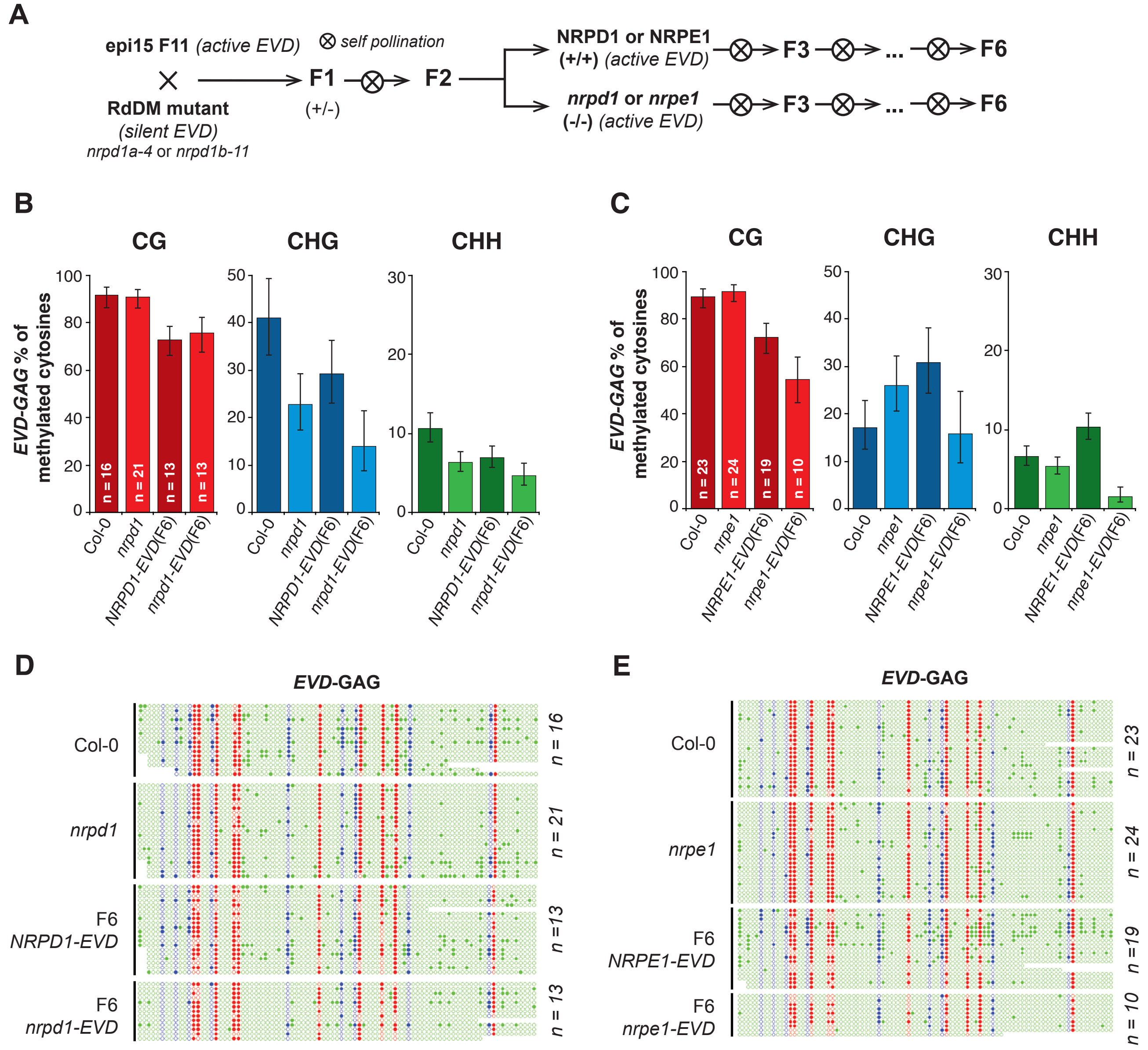


**F**

| | | Total sites | | | % of methylated C | | | | | Wilson Score Interval | | | | | |
|---|---|---|---|---|---|---|---|---|---|---|---|---|---|---|---|
| | | CG | CHG | CHH | CG | CHG | CHH | # of Clones | # of Exp | CG int- | CHG int- | CHH int- | CG int+ | CHG int+ | CHH int+ |
| **Controls - NRPD1** | Col-0 | 158 | 139 | 1069 | 0.9177 | 0.4100 | 0.1057 | 16 | 2 | 0.8643 | 0.3317 | 0.0887 | 0.9513 | 0.4931 | 0.1256 |
| | *nrpd* | 210 | 189 | 1461 | 0.9095 | 0.2275 | 0.0629 | 21 | 2 | 0.8630 | 0.1735 | 0.0516 | 0.9413 | 0.2923 | 0.0765 |
| **F6** | *NRPD1-EVD* | 199 | 178 | 1364 | 0.7286 | 0.2921 | 0.0689 | 20 | 2 | 0.6629 | 0.2303 | 0.0566 | 0.7856 | 0.3627 | 0.0836 |
| | *nrpd1-EVD* | 128 | 115 | 889 | 0.7578 | 0.1391 | 0.0461 | 13 | 2 | 0.6768 | 0.0875 | 0.0342 | 0.8238 | 0.2141 | 0.0619 |
| **Controls - NRPE1** | Col-0 | 228 | 205 | 1583 | 0.8947 | 0.1707 | 0.0656 | 23 | 2 | 0.8481 | 0.1254 | 0.0544 | 0.9282 | 0.2281 | 0.0789 |
| | *nrpe1* | 240 | 216 | 1676 | 0.9166 | 0.2592 | 0.0531 | 24 | 2 | 0.8747 | 0.2053 | 0.0434 | 0.9454 | 0.3215 | 0.0649 |
| **F6** | *NRPE1-EVD* | 188 | 169 | 1294 | 0.7234 | 0.3076 | 0.1027 | 19 | 2 | 0.6555 | 0.2429 | 0.0873 | 0.7824 | 0.3808 | 0.1204 |
| | *nrpe1-EVD* | 99 | 89 | 675 | 0.5454 | 0.1573 | 0.0148 | 10 | 2 | 0.4475 | 0.0961 | 0.0081 | 0.6400 | 0.2469 | 0.0270 |

**Supplementary Figure 6: BS-PRC analysis of *EVD-GAG* DNA methylation levels in RdDM mutants. A** Crossing scheme to generate *nrpd1*- and *nrpe1-EVD* lines. F2 plants were genotyped to select homozygous WT and mutant lines for each background and propagated through selfing until the F6 generation. **B, C** Bisulfite-PCR DNA methylation analysis at *EVD-GAG* sequences in the F6 generation of *NRPD1-EVD* (B) and *NRPE1-EVD* (C) lines, in both WT (darker shade) and mutant (lighter shade) backgrounds. Col-0, *nrpd1* and *nrpe1* were used as controls. Error bars represent 95% confidence Wilson score intervals. **D, E** Dot-plot representation of bisulfite-PCR sequencing data for the F6 generation of *NRPD1-EVD* (D) and *NRPE1-EVD* (E) lines, in both WT and mutant backgrounds, at *EVD-GAG* sequences. Filled circle represent methylated, empty circles unmethylated cytosines in the CG (red), CHG (blue) and CHH (green) context. Col-0 and *nrpd1* or *nrpe1* were used as control for the endogenous *EVD* copies. **F** Detailed Wilson score intervals of the bisulfite-PCR sequencing data DNA methylation analysis.

### SUPPLEMENTARY FIGURE 7

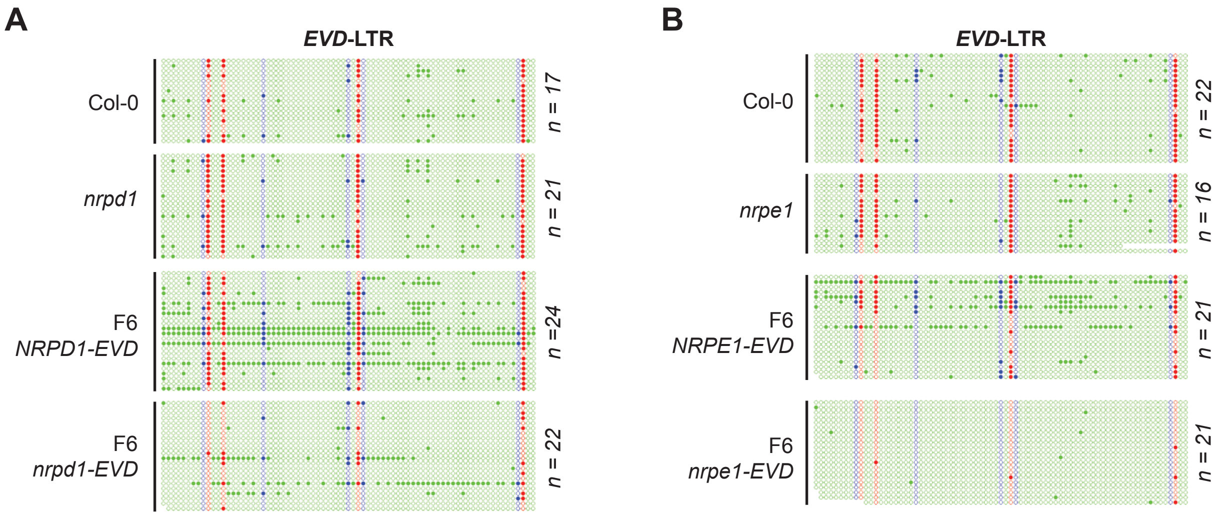


C

| | | Total sites | | | % of methylated C | | | | | Wilson Score Interval | | | | | |
|---|---|---|---|---|---|---|---|---|---|---|---|---|---|---|---|
| | | CG | CHG | CHH | CG | CHG | CHH | # of Clones | # of Exp | CG int- | CHG int- | CHH int- | CG int+ | CHG int+ | CHH int+ |
| **Controls - NRPD1** | Col-0 | 68 | 85 | 1121 | 0.6323 | 0.0705 | 0.0392 | 17 | 2 | 0.5135 | 0.0327 | 0.0293 | 0.7369 | 0.1454 | 0.0522 |
| | *nrpd* | 84 | 104 | 1386 | 0.8690 | 0.0865 | 0.0519 | 21 | 2 | 0.7805 | 0.0462 | 0.0414 | 0.9252 | 0.1563 | 0.0649 |
| **F6** | *NRPD1-EVD* | 96 | 120 | 1583 | 0.7916 | 0.3250 | 0.2154 | 24 | 2 | 0.6999 | 0.2478 | 0.1959 | 0.8608 | 0.4131 | 0.2363 |
| | *nrpd1-EVD* | 86 | 108 | 1440 | 0.2558 | 0.1203 | 0.0597 | 22 | 2 | 0.1754 | 0.0717 | 0.0486 | 0.3571 | 0.1950 | 0.0731 |
| **Controls - NRPE1** | Col-0 | 88 | 110 | 1449 | 0.8181 | 0.0909 | 0.0303 | 22 | 2 | 0.7248 | 0.0501 | 0.0226 | 0.8848 | 0.1593 | 0.0404 |
| | *nrpe1* | 63 | 79 | 1045 | 0.8730 | 0.0632 | 0.0430 | 16 | 2 | 0.7689 | 0.0273 | 0.0323 | 0.9342 | 0.1396 | 0.0571 |
| **F6** | *NRPE1-EVD* | 84 | 105 | 1384 | 0.4642 | 0.2380 | 0.1221 | 21 | 2 | 0.3615 | 0.1667 | 0.1059 | 0.5701 | 0.3278 | 0.1404 |
| | *nrpe1-EVD* | 83 | 104 | 1364 | 0.0602 | 0.0000 | 0.0058 | 21 | 2 | 0.0260 | 0.0000 | 0.0029 | 0.1333 | 0.0356 | 0.0114 |

**Supplementary Figure 7: BS-PRC analysis of *EVD-LTR* DNA methylation levels in RdDM mutants. A, B** Dot-plot representation of bisulfite-PCR sequencing data for the F6 generation of *NRPD1-* and *nrpd1-EVD* (A) or *NRPE1-* and *nrpe1-EVD* (B) lines, at *EVD-LTR* sequences. Filled circle represent methylated, empty circles unmethylated cytosines in the CG (red), CHG (blue) and CHH (green) context. Col-0 and *nrpd1* or *nrpe1* were used as control for the endogenous *EVD* copies. **C** Detailed Wilson score intervals of the bisulfite-PCR sequencing DNA methylation analysis.

### SUPPLEMENTARY FIGURE 8

**PCR/qPCR primers**

| Use | Name/Target | Sequence 5'>3' | Notes |
|---|---|---|---|
| Copy number/ Expresion levels | ACT2 F | GCACCCTGTTCTTCTTACCG | |
| Copy number/ Expresion levels | ACT2 R | AACCCTCGTAGATTGGCACA | |
| Copy number | EVD-GAG F | TTTGACCCGCGTGTTTGAAG | |
| Copy number | EVD-GAG R | AATCTTCGGGTCAAGCGTTC | |
| Copy number | EVD-IN F | CCGGAGAACAAAGAAGCAAGC | |
| Copy number | EVD-IN R | AATGTGCGGTTCTTGGTTGG | |
| Expression levels | shGAG F | GTTGGTTGCTACATCCACACCT | |
| Expression levels | shGAG R | TTTTCCCGTCTCAATATCCGGATT | |
| BiS-PCR | EVD-LTR F | GGATATGTATTATAAGAGAGAGTGGGTYGAATATATG | |
| BiS-PCR | EVD-LTR R | TTTATAARCATAAAAACATAATCTTATRCTCTAATACCATA | |
| BiS-PCR | EVD-3'GAG F | GTGTTTGAAGTGGAAGAAGGYGATTAAYGAATTAA | |
| BiS-PCR | EVD-3'GAG R | ATAACCCRACTTAACTTTRCTCCTCATAAATTTCTTAAA | |
| Genotyping | SAIL_LB3 | tagcatctgaatttcataaccaatctcgatacac | T-DNA BP primer for genotyping *rdr6-15* |
| Genotyping | rdr6-15 LP | TGAATCCATTCCTGAACAAGC | *mut rdr6-15*: BP x LP |
| Genotyping | rdr6-15 RP | CAATGCAACCTCATCTTGGATG | WT *RDR6*: LP x RP |
| Genotyping | SALK_LBa1 | TGGTTCACGTAGTGGGCCATCG | T-DNA BP for genotyping *nrpd1a-4* and *nrpd1b-11* |
| Genotyping | NRPD1B-11 LP | CAAAGTGGTGATGCATGGAGG | *mut nrpd1b-11*: BP x LP |
| Genotyping | NRPD1B-11 RP | ATGTAAATTTTGGAAGTCGGC | WT *NRPD1A-11*: LP x RP |
| Genotyping | NRPD1A-4 LP | GCACGGGTTCGAATACGGG | *mut nrpd1a-4*: BP x LP |
| Genotyping | NRPD1A-4 RP | GTATCTGACACCGCGGACTC | WT *NRPD1A-4*: LP x RP |

**Probes**

| Use | Target | Sequence 5'>3' | Notes |
|---|---|---|---|
| Northern blots | EVD-LTR F | ATGATGCTCGAGAGTGCGACAAGATCGATGTAGGT | PCR probe |
| Northern blots | EVD-LTR R | TACAATTCCGCATATTCTTTCATGGTATCAGAGCATA | PCR probe |
| Northern blots | EVD-GAG F | TAAGTCAAGAAGACTTAGAGTTTA | PCR probe |
| Northern blots | EVD-GAG R | AAGAAACTCATGAGGAGCAAAGT | PCR probe |
| Northern blots | siR1003 | ATGCCAAGTTTGGCCTCACGGTCT | Oligo probe |
| Northern blots | tasi255 | TACGCTATGTTGGACTTAGAA | Oligo probe |
| Northern blots | miR171 | GATATTGGCGCGGCTCAATCA | Oligo probe |
| Northern blots | U6 | AGGGGCCATGCTAATCTTCTC | Oligo probe |

**Supplementary Figure 8: Oligos used in this study**
